# Unified Network Spectroscopy: Inferring Reduced Biological Dynamics from Fluctuations under Structured Noise and Broken Ergodicity

**DOI:** 10.64898/2026.10.07.757233

**Authors:** Aleksandar Milinković, Lazar Stojanović

**Affiliations:** Union University - School of Computing, Belgrade, Serbia

**Keywords:** identifiability, ergodicity, anisotropy, Lyapunov equation, non-normality, multiscale

## Abstract

Mechanistic models of biological networks predict but cannot be calibrated at scale, while learned representations describe but expose no dynamical operator. Both lack an inverse step mapping data onto the few dynamical quantities the data determine. We develop Unified Network Spectroscopy (UNS) as that operator and establish when it is well posed. Reading fluctuations about a physiological setpoint through the Lyapunov relation of the linear noise approximation, UNS returns dominant timescales and frequencies, the subspace on which they act, and the projected noise. The inversion itself is established; our contribution is architectural. Three results govern its success. First, biochemical noise is generically anisotropic, and extrinsic modulation makes the dynamics non-normal. Assuming isotropy can invert mode rankings and destroy precision-matrix topology recovery; we give the exact, checkable condition under which recovery survives. Second, equal-time covariance identifies neither the dynamics nor their spectrum: the residual freedom is precisely the probability current that non-equilibrium living systems generically carry. Third, population snapshots replace time series only under ergodicity; heterogeneity biases inferred relaxation rates downward, making systems appear closer to bifurcation, whatever the sample size. Lagged covariance or per-cell velocity restores identifiability and turns noise structure into a recovered, falsifiable quantity. We provide a three-regime ergodicity taxonomy with diagnostics and show that the variational operator remains well posed when samples are fewer than variables. The output is a typed interface contract for hierarchical model composition, carrying its uncertainty, attainment and validity domain, with controllability inheritance as a falsification test.

## 1 Introduction

### 1.1 The Inverse Problem in Network Biology

A model of a biological network can be correct in every equation and still be unusable. This is an observation about where quantitative biology has stalled rather than a rhetorical flourish. Mechanistically faithful models exist at the molecular, cellular and tissue scales, and whole-cell simulation has demonstrated that an organism’s molecular processes can be represented in a single executable model [1–3]. What has not followed is the ability to use such models to decide what to do to a particular system.

The obstruction is that these are *forward* models: they map parameters to observations. Using them requires travelling in the opposite direction, and for models of this size that journey is not merely hard but ill-posed. A cellular model carries 10^2^ to 10^3^ kinetic parameters; the measurements available from a living system number in the tens. Fitting the former to the latter does not produce a calibrated model. It produces an arbitrary member of a large equivalence class, each element of which reproduces the data and each of which makes different predictions about intervention. Sensitivity analyses across systems-biology models show this directly: parameter sensitivities span many orders of magnitude, so most parameter combinations are unconstrained by any feasible experiment [4, 5].

The complementary paradigm fails for a mirror-image reason. Representation learning on biological data (e.g. foundation models) produces embeddings that predict phenotype and generalise across assays, but embedding is point in a coordinate system, not a dynamical system. It has no generator, no timescales, and no defined response to an input. One cannot ask a learned latent space what happens under perturbation, because the question requires an operator the representation does not contain.

#### The gap.

Rational intervention requires knowing how a system *responds*: its timescales, its stability margins, and the directions in state space along which a perturbation propagates. Forward mechanistic models contain this information but cannot be calibrated to recover it. Learned representations can be fitted to data but do not contain it. What is missing is an operator mapping measurements onto response characteristics directly, together with a statement of which response characteristics the measurements determine.

### 1.2 Forward and Inverse, in Concrete Terms

The words *forward* and *inverse* carry much of the argument, so it is worth fixing their meaning on an example small enough to check by hand.

Consider a single gene product present at copy number *x*, produced at a constant rate *α* and degraded in proportion to its own abundance at rate *βx*. The *forward* problem is: given (*α, β*), predict what will be observed. It has a complete answer. The mean settles at *ϕ*^∗^ = *α/β*; fluctuations about that mean have variance *α/β* as well; and a displacement from the mean decays at rate *β*, so that *R*(*τ* ) = (*α/β*) *e*^−*βτ*^ .

The *inverse* problem runs the other way: given observations, recover (*α, β*). The difficulty is already visible at this size.

A snapshot of a population (however many cells it contains, however precisely each is measured) determines only *α/β*. Doubling both parameters leaves every static observable unchanged. What separates them is the *timescale*, which is a property of how the system moves rather than of where it sits. A direct stochastic simulation with *α* = 60, *β* = 3 confirms the arithmetic: measured mean 20.02 against a predicted 20, variance 19.89 (Fano factor 0.99), autocorrelation decay rate 2.97 against a predicted 3.00, and hence *α* recovered as 59.4.

#### Inverse operator.

A map that runs opposite to a model: measurements in, dynamical quantities out. What makes one *usable* is a further requirement: that it returns only the quantities the measurements actually determine, together with an explicit statement of which quantities they do not. In the example above, an usable inverse operator given only a snapshot returns *α/β* and declares *α* and *β* individually unidentified. Given time-resolved data it would return both.

Everything that follows is this example with *n* interacting species instead of one. The parameter pair becomes the *n* × *n* Jacobian *J*; the variance becomes the covariance matrix *C*; the decay rate becomes the spectrum of *J*; and the observation that static data determine only a ratio becomes Theorem 5.2, which identifies exactly which *n*(*n* − 1)*/*2 combinations remain undetermined and why. The remedy is the same at both scales: resolve the measurement in time.

### 1.3 Scales, Interfaces, and the Composition Problem

Biological systems span an enormous range of scales, and no single model covers it. The standard response is to build a separate model at each level and couple them (a molecular model feeding a pathway model feeding a cell model, and so on up). The *interface* is the boundary between two adjacent models: the place where the finer model reports upward and receives context downward (Figure 1).

**Figure 1.**
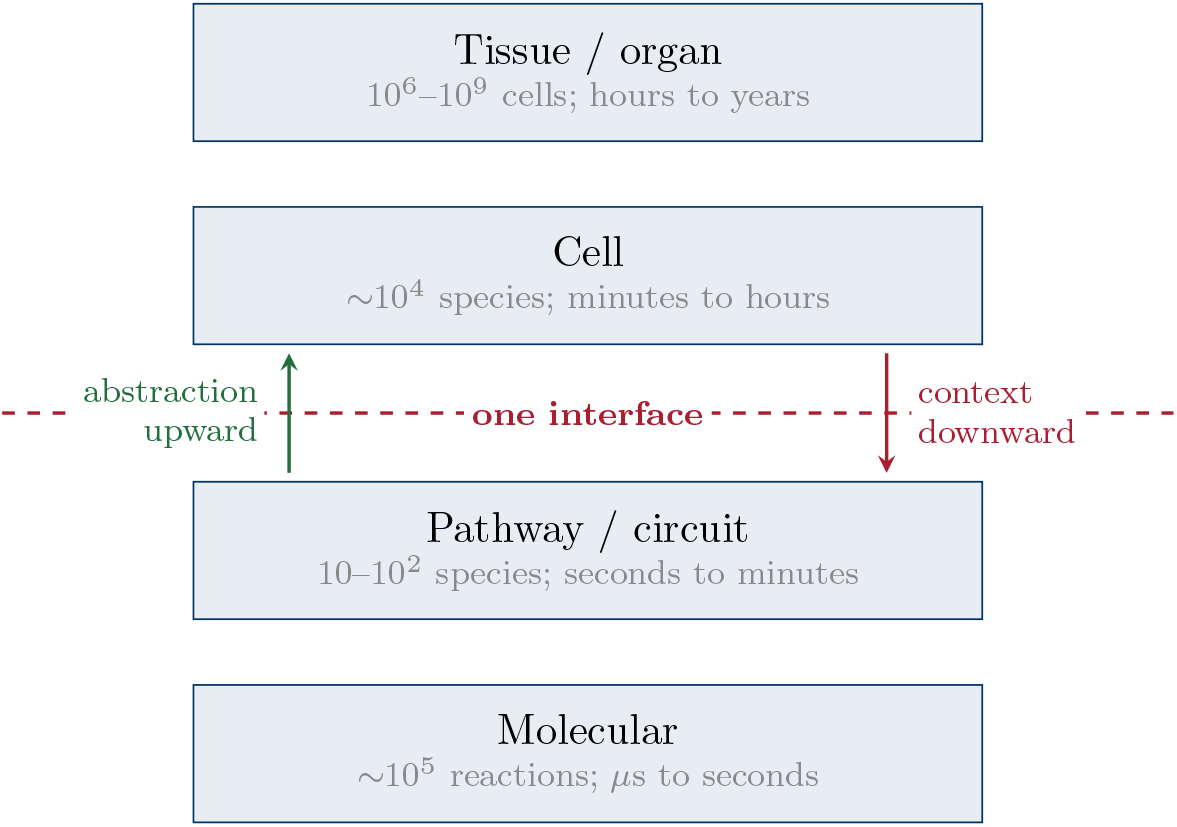
A hierarchy of models and the interface between two of its levels. Only a small, declared set of quantities may cross: upward, a compressed dynamical description; downward, the operating point and constraints that the coarser level imposes. The design question this article addresses is what that upward set should be.

The point of an interface is that the coarser model does not need everything the finer one knows. A cell-level model does not need each of the 10^4^ molecular concentrations beneath it. It needs to know how fast the pathway responds, whether that response is stable, in which combinations of species the response appears, and how noisy the output is — a handful of numbers. Deciding *which* handful is the interface design problem, and it has two standard answers that both fail.

#### Pass everything upward

Then nothing has been compressed, and parameter counts multiply from one level to the next until the composite model has more free quantities than any experiment can constrain.

#### Pass fitted mechanistic parameters upward

Then the numbers being passed are, by the argument of Section 1.2, not determined by the data. They are one arbitrary choice from a large equivalent set. Each level inherits and compounds the arbitrariness of the level below.

The requirements of Section 1.5 are what remains once both answers are rejected.

### 1.4 What a Contract Specifies

The word *contract* is borrowed from software and systems engineering, where it has a precise meaning. A contract attached to a component states three things: a **guarantee**, being what the component promises about its own behaviour; an **assumption**, being what it requires of its environment for that promise to hold; and the **conditions of validity** under which the promise applies at all. A sorting routine guarantees ordered output, assumes the comparison it is handed is a total order, and offers its performance guarantee only while the input fits in memory. The value of stating this explicitly is that a system assembled from such components can be checked at the joins, without reopening the components.

Transposed to the present setting the guarantee is spectral: this subsystem relaxes with these timescales, along these directions, with this much noise. The assumption is the operating regime: near this setpoint, over this observation window, with data of this kind. The conditions of validity are recorded explicitly rather than left to the reader’s judgement.

Two features distinguish a contract from an ordinary reduced model. First, the obligations run in *both* directions: a coarser model built on top must not contradict the guarantee, and any finer refinement below must reproduce it. A reduced model is something one uses; a contract is something one can violate. Second, because it can be violated, it can be checked. Which is what makes composition auditable rather than merely convenient.

The qualifier *typed* means that only objects of the declared kind may cross the interface. Presenting anything else (a fitted rate constant, a raw concentration, an embedding coordinate) is a violation of the specification rather than a variation on it.

### 1.5 Requirements on an Interface Quantity

Suppose models are to be composed hierarchically, each scale exporting a compressed description upward. What should cross that boundary? The following requirements eliminate almost every candidate.

#### R1. Identifiability

The exported quantity must be determinable from data that can realistically be collected. Exporting a quantity the data cannot resolve creates the appearance of a calibrated interface while leaving its value arbitrary.

#### R2. Compression

The exported quantity must be far lower-dimensional than the state it summarises, or the hierarchy accomplishes nothing.

#### R3. Dynamical content

It must carry information about *evolution*, not merely about static structure or variance. A correlation matrix, a cluster label, or a set of principal components describes how a system is configured; none says how fast it relaxes or whether it is near instability.

#### R4. Actionability

It must be of a type that control synthesis consumes: a generator, an input map, and a noise model.

Table 2 scores the candidates. Only a spectral description of the local generator satisfies all four, and its identifiability entry is marked conditional, a qualification that Sections 3–5 exist to make precise.

**Table 1:** What each measurement of a single birth–death process determines. The mean and the variance are numerically different quantities that carry the *same* information; only a time-resolved measurement separates production from degradation.

| Measurement | Value | Information gained |
| --- | --- | --- |
| Mean, $\phi^*$ | $\alpha/\beta$ | the ratio $\alpha/\beta$ |
| Variance, $C$ | $\alpha/\beta$ | nothing new — the same ratio |
| Autocorrelation decay rate | $\beta$ | $\beta$, and hence $\alpha = \beta\phi^*$ |

**Table 2:** Candidate interface quantities against the requirements of Section 1.5.

| Candidate | R1 Ident. | R2 Compr. | R3 Dynam. | R4 Action. |
| --- | --- | --- | --- | --- |
| Raw state variables | yes | **no** | partial | partial |
| Mechanistic kinetic parameters | **no** | **no** | yes | yes |
| Learned embedding $z = \phi(x)$ | yes | yes | **no** | **no** |
| Correlation / covariance matrix | yes | **no** | **no** | **no** |
| Principal components | yes | yes | **no** | partial |
| Koopman eigenfunctions | partial | partial | yes | yes |
| **Spectral triple** ($T_r, U_r, Q_r$) | **cond.** | **yes** | **yes** | **yes** |

**Table 3:** Diffusion matrix computed from (4) for a three-species network (Ω = 1). Left: independent birth–death kinetics with transcription rates (1000, 50, 5). The matrix is diagonal but its entries span a factor of 200. Right: the same network plus one bimolecular reaction *X*_1_ +*X*_2_ → *X*_3_ with propensity 30. Off-diagonal entries appear with *both* signs, as Proposition 3.1(iii) predicts.

| Birth–death only | With $X_1 + X_2 \rightarrow X_3$ |
| --- | --- |
| $\begin{pmatrix} 2000 & 0 & 0 \\ 0 & 100 & 0 \\ 0 & 0 & 10 \end{pmatrix}$ | $\begin{pmatrix} 2030 & 30 & -30 \\ 30 & 130 & -30 \\ -30 & -30 & 40 \end{pmatrix}$ |

**Table 4:** Two modes with *λ*_1_ = −0.05 and *λ*_2_ = −5.0, so that mode 1 is 100× slower. With comparable noise power the slow mode dominates the covariance, as expected. With modal noise power differing by a factor of 500 — well within the range spanned by transcription rates in a single cell — the *fast* mode dominates, and any procedure that ranks modes by covariance magnitude retains the wrong one.

| Modal noise power ($\tilde{Q}_{11}, \tilde{Q}_{22}$) | Modal variance | Dominant mode | Correct? |
| --- | --- | --- | --- |
| (1, 1) | (10.0, 0.1) | mode 1 (slow) | yes |
| (1, 500) | (10.0, 50.0) | mode 2 (fast) | **ranking inverted** |

**Table 5:** Magnitude of the bias of Corollary 4.2 for *k*_true_ = 0.8, *q* = 1, so that *C*_w_ = 0.625. Static heterogeneity comparable to within-cell variance already inflates the apparent relaxation time by a factor of 2.6.

| $\text{Var}_\theta(x^*)$ | $C_w$ | $C_{cs}$ | $\hat{k}$ | $\hat{\tau}/\tau_{\text{true}}$ |
| --- | --- | --- | --- | --- |
| 0.00 | 0.625 | 0.625 | 0.800 | 1.00 |
| 0.25 | 0.625 | 0.875 | 0.571 | 1.40 |
| 1.00 | 0.625 | 1.625 | 0.308 | 2.60 |
| 4.00 | 0.625 | 4.625 | 0.108 | 7.40 |

The spectral triple comprises the reduced generator *T*_*r*_ (timescales, frequencies, stability margins), an orthonormal basis *U*_*r*_ for the retained invariant subspace, and the modal noise covariance *Q*_*r*_; it is constructed in Section 6.3. Because biological generators are not normal, the description is stated on the real Schur form rather than on eigenvectors: the eigen-decomposition is what one interprets, but it is not always defined, and Remark 6.3 gives the dictionary between the two.

### 1.6 Why Spectra, and What This Article Adds

Three observations motivate a spectral interface. Eigenvalues of the linearised generator are what control synthesis needs: they set relaxation timescales *τ*_*i*_ = −1*/*Re(*λ*_*i*_), oscillation frequencies *ω*_*i*_ = Im(*λ*_*i*_), and distance to instability. Fluctuation statistics preferentially reveal them, because modal variance scales inversely with modal damping. And they are constrained by network structure, which supplies falsification tests unavailable to purely statistical methods.

Each of these three claims requires qualification once realistic noise is admitted, and supplying those qualifications is the purpose of this article. The second claim in particular (that fluctuations reveal slow modes) holds only up to a weighting by noise power, and when that weighting spans orders of magnitude, as it does in gene expression, the ranking of modes by covariance can invert entirely (Section 3.5).

We assume throughout that the reader is interested in the operator as a practical inference device. Accordingly, every structural claim is accompanied either by a proof or by a numerical demonstration, and Section 15 states the conditions under which the operator should not be applied at all.

### 1.7 Scope

UNS is an inference operator: a map from fluctuation statistics to a reduced dynamical description, with uncertainty estimates and an explicit declaration of what was not identifiable. It is not a modelling framework, a simulator, or a substitute for mechanistic knowledge. It does not assume biological systems are linear; it exploits the local linear structure that homeostatic feedback actively maintains near setpoints (Section 8), and reports the domain over which that structure is supported by data.

It is not novel as a technique. Section 2 sets out the inheritance in detail. What is claimed as new is the use of the resulting spectral triple as a typed interface contract with enforced bidirectional consistency (Section 14), together with the noise-structure and ergodicity analyses that determine when the contract terms are estimable at all.

### 1.8 A Note on the Name

The term is meant literally, and the analogy is closer than it may first appear. Spectroscopy does not require driving a system. Newton’s decomposition of sunlight by prism [6] and the spectral chemical analysis founded by Kirchhoff and Bunsen [7] are entirely passive, as is all astronomical spectroscopy. What defines the method is the *resolution of a signal into its component frequencies* in order to infer the structure of the system that produced it. Emission spectroscopy does this with light a source radiates spontaneously; UNS does it with the fluctuations a biological network produces spontaneously.

The direct methodological ancestors are the fluctuation-correlation spectroscopies. Fluorescence correlation spectroscopy [8–11] extracts diffusion coefficients and reaction rate constants from the autocorrelation of spontaneous concentration fluctuations in a femtolitre volume; dynamic light scattering, also known as photon correlation spectroscopy [12], does the same for particle dynamics; and electronic noise spectroscopy infers dissipative properties from thermal voltage fluctuations [13, 14]. All are passive. UNS is the network-scale member of this family.

Taking the analogy seriously sharpens the central technical point of this article. The stationary covariance is a *frequency-integrated* quantity:

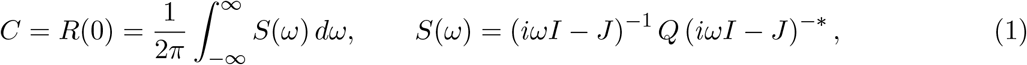

that is, the total power with all frequency information discarded. Measuring *C* alone is therefore not spectroscopy at all but *photometry* : it is the analogue of recording a star’s total brightness with no prism in the beam. The lagged covariance *R*(*τ* ), equivalently the power spectral density *S*(*ω*), is precisely what disperses that power across frequencies.

Section 5.2 shows that the generator is not recoverable from *C* alone. Read optically, this is the unsurprising statement that a source’s line structure cannot be inferred from its integrated brightness, and the remedy is correspondingly simple: put the prism in.

One caveat survives, and it is narrower than the naming question. Inferring a system’s *response to perturbation* from its spontaneous fluctuations is a further step, licensed by the fluctuation–dissipation relationship [15, 16], which holds at thermal equilibrium and fails in the non-equilibrium steady states that living systems maintain. Emission spectroscopy faces the same limitation in its own domain: line positions give level spacings, but oscillator strengths (the response) require more than the passive spectrum. Section 5.2 identifies exactly which component of the dynamics is lost as a result, and Section 5.4 recovers it.

### 1.9 Roadmap

Section 2 sets out the prior art, since the operator developed here has been invented independently in three fields and the contribution claimed is one of composition rather than of technique. Sections 3 and 4 are the two prerequisites that determine whether the inference is well posed at all: the first establishes what the driving noise of a biochemical network actually looks like and what goes wrong when it is assumed isotropic; the second establishes when a population snapshot may stand in for a time course. Section 5 contains the central identifiability result and its remedies, Section 6 defines the operator, and Section 7 treats uncertainty and the amount of data required. Sections 8–11 concern the biology: why physiological circuits admit a local linear description at all, what network structure implies about the inferred spectrum, and three worked circuits. Sections 12 and the following one relax stationarity and linearity. Section 14 returns to the interface contract with the machinery in place, and Section 15 states the conditions under which none of this should be applied.

A reader interested only in whether the method applies to a particular dataset may find Table 6, Protocols 5.7 and 5.8, and the limitations of Section 15 sufficient.

**Table 6:** Regimes for the substitution of ensemble for temporal statistics. *T*_obs_ is the observation window; *τ*_slow_ the slowest retained mode; *T*_erg_ the time to explore the accessible state space.

| Regime | Condition | Cross-sectional data | Required action |
| --- | --- | --- | --- |
| **E**<br>ergodic | $\tau_{\text{slow}} \ll T_{\text{obs}}$, population homogeneous | Estimates $C$ consistently; gives no lag information | Use for $C$; obtain $J$ from velocity statistics (§4.7) or from paired longitudinal data |
| **H**<br>heterogeneous | $\text{Cov}_\theta(x^*)$ non-negligible | Biased by Theorem 4.1; relaxation appears too slow | Stratify by covariates; use dual-reporter or allele-paired designs to estimate the split; or use longitudinal data |
| **N**<br>non-ergodic | $\tau_{\text{slow}} \gtrsim T_{\text{obs}}$ or $T_{\text{erg}} \rightarrow \infty$ | Measures a distribution over frozen states, not a dynamical covariance | Exclude affected modes from the exported description; model separately as a hidden discrete state |

**Table 7:**
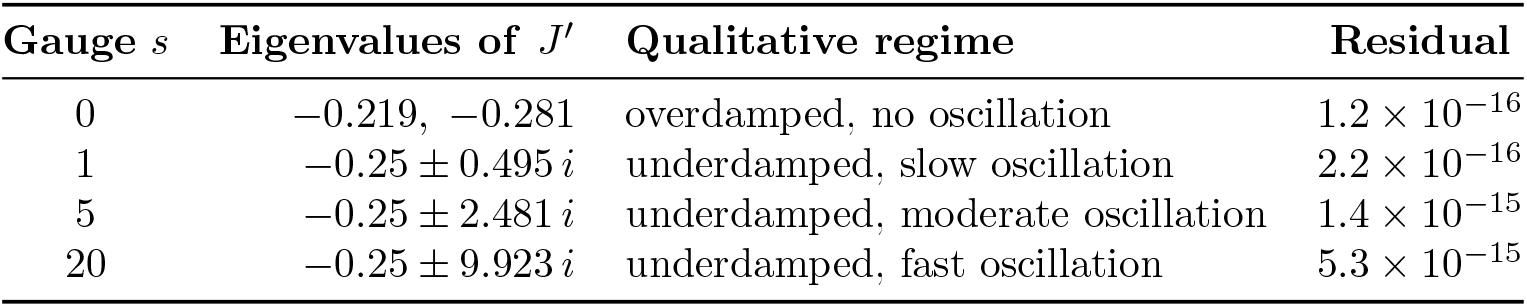
Four generators consistent with *identical C* and *Q*, obtained by varying the gauge 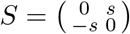 in Theorem 5.2, for the damped oscillator (39) with *ω* = 2, *γ* = 0.5, *Q* = *I*. The residual column confirms each satisfies (17) to machine precision. Recovered dynamics range from purely overdamped to oscillation at forty times the true frequency.

## 2 Prior Art

The operator developed here has been invented independently at least three times, in fields that do not read each other’s journals. Setting out that history is not a formality: an architectural contribution consists in the disciplined composition of existing components, and such a claim can only be assessed if the components are correctly identified.

### Stochastic kinetics and the linear noise approximation

The mathematical foundation is van Kampen’s system size expansion [17], which yields the deterministic rate equations at leading order and a linear stochastic differential equation for the fluctuations at next order. Its use for biochemical networks was developed by Elf and Ehrenberg [18] and has since been systematised in [19–21]. The exact underlying description is the chemical master equation, simulated by the Gillespie algorithm [22, 23] or its accelerated variants.

### Fluctuation-correlation spectroscopy

The idea of extracting kinetic constants from spontaneous fluctuations rather than from imposed perturbations predates its network-scale form by half a century. Fluorescence correlation spectroscopy, introduced by Magde, Elson and Webb to measure a chemical relaxation rate from equilibrium concentration fluctuations [8, 9], remains a standard tool for diffusion and binding kinetics in living cells [10, 11]; dynamic light scattering applies the same principle to particle motion [12]; and the measurement of dissipative properties from thermal electrical noise [13, 14] is the original instance. What these methods do for one or two degrees of freedom, UNS attempts for a coupled network. The essential methodological inheritance is that all of them analyse the *time-resolved* correlation function rather than the variance alone, a point whose necessity Section 5.2 establishes formally.

### The inverse Jacobian in metabolomics

The most direct antecedent is a continuous line of work in metabolomics. Steuer, Kurths, Fiehn and Weckwerth [24] showed that correlations among metabolite concentrations carry information about the Jacobian of the reaction network, related through precisely the Lyapunov equation used here. Their formulation is identical to ours up to noise normalisation. The programme has continued for two decades: differential Jacobian estimation [25], identifiability of Jacobian alterations between conditions [26], a catalogue of practical obstructions [27], integration with genome-scale reconstruction [28], and most recently the combination of network structure with fluctuation data including a non-diagonal noise matrix derived from stoichiometry [29]. The last of these anticipates the information-fusion scheme of Section 10.5, and that section should be read as an adaptation rather than an independent invention.

### Effective connectivity in neuroscience

The same operator appears in whole-brain modelling as multivariate Ornstein–Uhlenbeck effective connectivity [30, 31]. Significantly for the present article, that work uses *both* zero-lag and lagged covariance, precisely because the zero-lag covariance alone is insufficient (the limits of the equal-time-only case having been examined explicitly [32]). Neuroscience also supplies the vocabulary: the distinction drawn in Section 10.3 between the network of possible interactions and the network of dynamically active influence is the long-established structural/effective connectivity distinction [33, 34].

### Graph learning from signals

A third community reached the same relationship from signal processing: a signal diffusing over a graph under isotropic forcing has a stationary covariance that is a matrix function of the graph Laplacian, so the precision matrix recovers the Laplacian up to a diagonal shift [35–39], surveyed in [40, 41]. The statistical ancestor is Gaussian graphical modelling [42], and the biological analogue is the separation of direct from indirect dependencies [43–45]. Section 10.1 restates this correspondence, and then shows that its isotropy hypothesis fails for biological noise, with consequences that have not, to our knowledge, been quantified.

### Noise decomposition in single cells

The partition of cellular variability into intrinsic and extrinsic components was formalised by Swain, Elowitz and Siggia [46] and validated by dual-reporter assays [47, 48], with theoretical development by Paulsson [49, 50] and a dynamic generalisation by Hilfinger and Paulsson [51]. Section 3.3 builds directly on this decomposition, which we argue is the same object as the ergodicity question of Section 4 viewed from a different angle.

### Model reduction

When a system is driven by white noise with covariance *Q* = *BB*^⊤^, the stationary covariance *C is* the controllability Gramian. Selecting modes by the dominant eigenvectors of *C* is therefore controllability-Gramian-only truncation, which is balanced proper orthogonal decomposition without the observability half [52–55]. UNS is not a fundamentally different reduction; it is the reduction one can perform with the Gramian that passive data deliver.

#### Statement of the novelty claim.

*Not claimed as novel:* the Lyapunov inversion; spectral reduction of the inferred generator; the precision-matrix/Laplacian correspondence; the intrinsic–extrinsic decomposition; the structural/effective topology distinction; topology-regularised inference; sliding-window spectral tracking; piecewise-linear regime models; eigenvalue-based early warning; spectral regularisation of learned latent dynamics.

*Claimed as new:* (i) the exact condition under which the precision-matrix/Laplacian correspondence survives anisotropic forcing, which is weaker than isotropy, together with the quantitative consequences of violating it for mode ranking and for topology recovery (Sections 3.5 and 10.1); (ii) the three-regime ergodicity taxonomy with decision diagnostics, and the systematic direction of the cross-sectional bias (Section 4); (iii) joint recovery of (*J, Q*) from velocity statistics, with positive semidefiniteness of the recovered *Q* as a misspecification test (Section 5.5); (iv) the spectral triple as a typed interface contract, with controllability inheritance as a falsification criterion for abstraction, and with attainment against a design-matched ceiling as a required qualifier alongside uncertainty (Section 14); and (v) the separation of structural from sampling rank deficiency in the covariance estimate, with the demonstration that the standard repairs fail in opposite directions (Section 7.3).

## 3 The Structure of Biological Noise

Most treatments of fluctuation-based inference assume, explicitly or tacitly, that the driving noise is isotropic: *Q* = *qI*. This section shows that the assumption is never even approximately correct for a biochemical network, characterises what *Q* actually looks like, and quantifies what goes wrong when the assumption is made anyway.

### 3.1 From the Chemical Master Equation to the Linear Noise Approximation

Consider a well-mixed compartment of size Ω containing *n* species subject to *R* reactions with stoichiometry matrix *S* ∈ ℤ^*n×R*^ and propensities *a*_*r*_(·). The exact description is the chemical master equation [17, 22]. Writing the copy numbers as

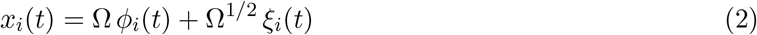

and expanding in Ω^−1*/*2^ yields the macroscopic rate equations 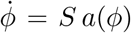 at leading order and, at next order, a linear stochastic differential equation for the fluctuations [17, 18]:

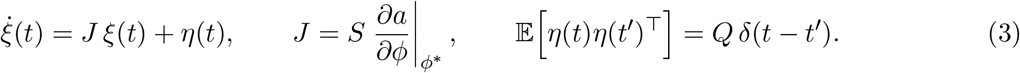

The essential point, and the one usually passed over, is that the same expansion that produces *J* also produces *Q*:

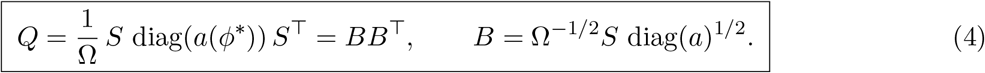

*Q* is not a free parameter to be assumed. It is a determinate function of the network’s stoichiometry and its operating point.

### 3.2 Why *Q* Is Generically Anisotropic

Equation (4) has four structural consequences.

#### Proposition 3.1

(Structure of the diffusion matrix). *Let Q be given by* (4) *with a*_*r*_(*ϕ*^∗^) *>* 0 *for all r*.

*Then*

i. *Q* ⪰ 0, *and* rank(*Q*) ≤ min(*n, R*);
ii. *if* ℓ ∈ ℝ^*n*^ *satisfies* ℓ^⊤^*S* = 0 *(a conservation law), then Q*ℓ = 0, *so Q is singular;*
iii. *Q*_*ij*_ = Ω^−1^ ∑_*r*_ *S*_*ir*_*S*_*jr*_*a*_*r*_, *which is nonzero whenever some reaction alters both species i and j, and whose sign is that of the stoichiometric product — negative for a conversion X*_*i*_ → *X*_*j*_, *positive for co-consumption or co-production;*
iv. *Q* = *qI requires S* diag(*a*)*S*^⊤^ ∝ *I; in particular every reaction must alter exactly one species with unit stoichiometry, and all propensities at the operating point must be equal*.

*Proof*. (i) *Q* = *BB*^⊤^ is a Gram matrix, and rank(*BB*^⊤^) = rank(*B*) ≤ min(*n, R*). (ii) ℓ^⊤^*S* = 0 gives *Q*ℓ = Ω^−1^*S* diag(*a*)*S*^⊤^ℓ = 0. (iii) and (iv) are immediate from the definition. □

Condition (iv) is worth dwelling on, and it is worth stating carefully, because two claims are easily conflated here and only the weaker one is true. Even the simplest case — *n* independent genes each with birth–death kinetics ∅ → *X*_*i*_ at rate *α*_*i*_ and *X*_*i*_ → ∅ at rate *β*_*i*_*x*_*i*_ — gives *Q* = Ω^−1^ diag(2*α*_*i*_) at steady state, which is diagonal but proportional to the identity only if all transcription rates coincide. Since expression levels in a single cell span four to five orders of magnitude [56, 57], so does *Q*.

#### Remark 3.2

(Exact isotropy is possible but non-generic). Condition (iv) constrains the product *S* diag(*a*)*S*^⊤^, not the reaction list, so it can be met by cancellation between reactions rather than by every reaction altering a single species. Balanced stoichiometries admissible at steady state exist for which *Q*_int_ ∝ *I exactly*, even though every reaction alters two species. The correct statement is therefore not that biochemical diffusion matrices are never isotropic but that they are **generically anisotropic**, and the argument is one of measure rather than of exclusion: within the cone of admissible steady-state fluxes, the isotropic set is a lower-dimensional subset, so isotropy is attained only on a null set and approached only for flux vectors close to it.

The distinction matters in two directions. It weakens any claim that isotropy is *impossible*, which the results below do not need. And it strengthens the operational recommendation, which is not that isotropy is forbidden but that it is a substantive hypothesis about the operating point: it may be adopted where a stoichiometry and a flux distribution are available to justify it, and must not be adopted as a default. The same asymmetry recurs in Section 10.1, where a proper anisotropic subfamily preserves the precision-matrix correspondence that generic anisotropy destroys.

### 3.3 Intrinsic and Extrinsic Contributions

Equation (4) describes *intrinsic* noise: the randomness of individual reaction events within one cell. Cellular variability also has an *extrinsic* component arising from fluctuations in global modulators (e.g. ribosome and polymerase abundance, cell volume, cell-cycle phase, metabolic state, etc.) which affect many reactions simultaneously [46–48]. Extrinsic noise is not merely additional; it has an entirely different structure.

Let *θ*(*t*) = *θ*^∗^ + *δθ*(*t*) ∈ ℝ^*p*^ collect the global modulators and write 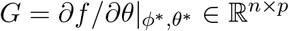 for the sensitivity of the drift to them. Then

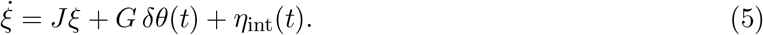

If *δθ* were white with covariance Σ_*θ*_, the total diffusion matrix would be

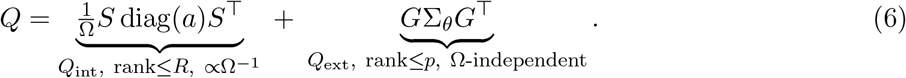

Three properties of *Q*_ext_ matter.

#### It is low rank and highly anisotropic

The number of independent global modulators is small: volume, ribosome pool, cell-cycle phase, energy charge, perhaps a handful more. So *Q*_ext_ has rank at most *p* ≪ *n* and concentrates all its power in the column space of *G*, i.e. along the directions in which the network is most sensitive to global parameters. Isotropy is not approached even asymptotically.

#### It does not vanish with system size

*Q*_int_ scales as Ω^−1^ while *Q*_ext_ does not. In high-copy-number systems extrinsic noise therefore dominates, which is the observed pattern: highly expressed genes have variability dominated by extrinsic sources, weakly expressed genes by intrinsic bursting [56–58].

#### It is coloured, not white

Global modulators fluctuate on the timescale of the cell cycle (hours) which is comparable to or slower than the network dynamics of interest. Modelling them as white noise is therefore wrong. The correct treatment introduces an explicit generator for the modulators, 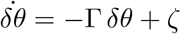, and augments the state:

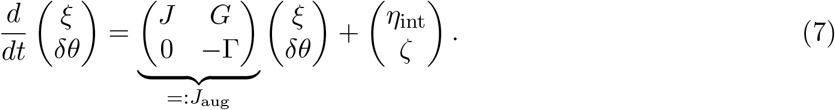

##### Remark 3.3

(Extrinsic noise manufactures non-normality). *J*_aug_ is block upper triangular with *G* ≠ 0 and is therefore non-normal whenever *J* and −Γ do not commute, which is generic. Even a network whose intrinsic generator is normal acquires a non-normal augmented generator through extrinsic coupling. Non-normality in biological fluctuation data is thus not an exotic possibility but a structural consequence of the way cells are embedded in a fluctuating global environment. Section 3.6 develops the consequences.

##### Remark 3.4

(Extrinsic modes contaminate the inferred spectrum). spec(*J*_aug_) = spec(*J*) ∪ spec(−Γ). An analysis that ignores the augmentation will attribute the modulator relaxation rates spec(−Γ) to the network under study. Since cell-cycle and metabolic timescales are slow, they appear among the *dominant* modes and are precisely the ones a reduced-order description retains. Reporting a cell-cycle eigenvalue as a property of a signalling network is a failure mode we expect to be common, and Section 10.4 gives a test for it.

### 3.4 Measurement Noise is Assay-Specific and Rarely Additive

A third contribution enters at the point of observation. What is analysed is not *x* but an assay-dependent readout, and the statistics of the readout differ qualitatively between platforms.

In droplet-based single-cell RNA sequencing, transcripts are captured with low and variable efficiency, so the observed count is approximately a binomial thinning of the true count. This is *multiplicative and count-dependent*, not additive Gaussian. Careful analyses show that the excess zeros in such data are explained by the sampling model itself rather than by a separate zero-inflation process [59, 60]. Modelling it as an additive *Q*_tech_ is a misspecification, and it biases the Fano factor upward at low expression. Spike-in based methods [61, 62] and hierarchical models [63] exist precisely to separate this contribution.

Flow cytometry and fluorescence microscopy have different characteristics again: autofluorescence contributes an additive floor, detector shot noise contributes a variance proportional to signal, and reporter maturation acts as a low-pass filter that colours the observed noise [58]. Live-cell imaging with destabilised reporters introduces a further timescale through reporter turnover.

The practical consequence is that

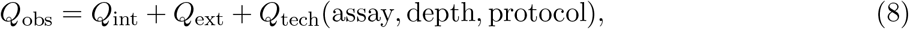

and the third term is not transferable between datasets. This is the formal content of the observation that the noise model depends on the concrete dataset: it does and a method that hard-codes *Q* cannot be applied across platforms. Section 5.5 removes the need to hard-code it.

### 3.5 The Ranking of Modes Can Invert

We now quantify the first consequence of non-isotropy.

Transform (3) to the eigenbasis of *J*: with 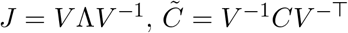 and 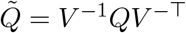, the Lyapunov equation gives the exact modal relation

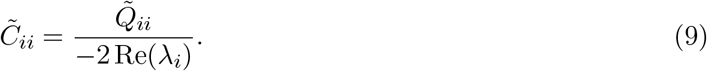

The usual reading of (9) is that slow modes dominate the covariance. That reading is correct only if 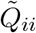 is comparable across modes. What (9) actually says is that modes are ranked by the ratio of noise power to damping, and mode *i* dominates mode *j* precisely when

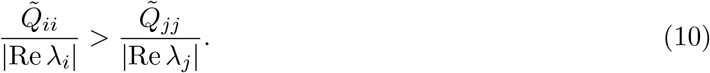

#### Algorithmic consequence.

Modes must be selected by estimated |Re *λ*_*i*_|, not by covariance magnitude. Since eigenvalues require lag or velocity information (Section 5.4), covariance-only mode selection is unsound whenever the noise is anisotropic, that is, always. Eigendecomposition of *Ĉ* remains useful for identifying a dominant *subspace* for dimension reduction, but not for ordering modes by timescale.

### 3.6 Non-Normality

A matrix is normal if *JJ*^⊤^ = *J*^⊤^*J*, equivalently if its eigenvectors can be chosen orthogonal. Biological generators are typically not normal: feedforward cascades, signalling chains, and (by Remark 3.3) extrinsic coupling all produce directional influence structures. The consequences are characterised in the matrix-analysis literature [64, 65] and documented for biological networks [66–69].

Three consequences matter here.

#### Transient amplification

A perturbation may grow by orders of magnitude before decaying, even with every eigenvalue in the left half plane; sup_*t*≥0_ ∥*e*^*Jt*^∥ is not bounded by any function of the spectrum alone. A controller specification derived only from spec(*T*_*r*_) can be met while the closed-loop system shows large excursions.

#### Spectral fragility

By the Bauer–Fike theorem, if *J* = *V* Λ*V* ^−1^ is perturbed by *E*, every eigenvalue *µ* of *J* + *E* satisfies

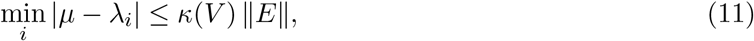

where *κ*(*V* ) is the eigenvector condition number. For normal *J, κ*(*V* ) = 1; for strongly non-normal *J* it may be enormous, and inferred eigenvalues are correspondingly unreliable.

#### Covariance eigenvectors are not dynamical modes

Relation (9) holds in the eigenbasis of *J*, which coincides with that of *C* only when *J* is normal *and Q* ∝ *I*. Neither holds. For a randomly generated stable non-normal *J* with *Q* = *I* in dimension 5, the descending eigenvalues of *C* and the sorted values of 1*/*|Re *λ*_*i*_| are

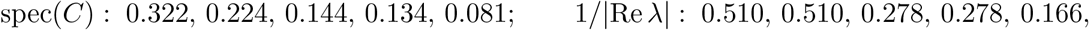

with ratios 0.630, 0.439, 0.518, 0.483, 0.490, not constant. Reading timescales off the covariance spectrum is therefore invalid.

The response adopted here is to report 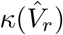 alongside 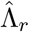 as part of the exported description, to use pseudospectral rather than spectral bounds where robustness matters, and to use the biorthogonal projection of Section 6.3. Pseudospectral analysis has been applied to the hypothalamic–pituitary– adrenal axis [70] one of the circuits considered in Section 11) with the conclusion that the eigenvalue picture alone is insufficient there.

## 4 When Ensemble Statistics Replace Temporal Statistics

The operator requires the stationary covariance *C* and, as Section 5.4 will show, the lagged covariance *R*(*τ* ). How these are estimated depends on what kind of data one has, and the two available kinds are not interchangeable.

### 4.1 Two Kinds of Data

*Cross-sectional* (ensemble, snapshot) data measure many individuals once. Single-cell RNA sequencing, flow cytometry, mass cytometry, spatial transcriptomics, and any destructive assay are of this kind. They deliver an empirical distribution over the population at one instant, hence an estimate of an ensemble covariance.

*Longitudinal* data follow one individual through time. Live-cell imaging with fluorescent or FRET reporters, microfluidic mother-machine experiments, and continuous physiological monitoring are of this kind. They deliver *Ĉ* and 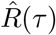 from a single trajectory, but at low throughput.

Cross-sectional data are abundant and cheap; longitudinal data are scarce and expensive. There is therefore strong practical pressure to substitute the former for the latter. The question is when this is legitimate.

### 4.2 The Ergodic Hypothesis, Stated Precisely

Substituting ensemble for temporal statistics requires three distinct conditions, which are frequently conflated.

#### E1. Stationarity

Each trajectory is statistically stationary over the observation window.

#### E2. Mixing

The process explores its state space on a timescale short compared with observation. For an Ornstein–Uhlenbeck process with Hurwitz *J* this holds automatically, with mixing time *τ*_mix_ = 1*/*|Re *λ*_slow_|; it fails when a mode is slower than the window.

#### E3. Population homogeneity

Every member of the ensemble is governed by the *same* parameters (*J, Q, x*^∗^).

E1 and E2 are properties of the dynamics and are shared with the longitudinal case. E3 is specific to the ensemble substitution, and it is the one biology violates. It is also the one most often left unstated.

### 4.3 The Cross-Sectional Covariance is Biased

#### Theorem 4.1

(Decomposition of the cross-sectional covariance). *Let a population be indexed by a latent parameter θ* ∼ *π, and let a cell with parameter θ have deterministic steady state x*^∗^(*θ*) *and within-cell stationary covariance C*_w_(*θ*) *solving J*(*θ*)*C*_w_ + *C*_w_*J*(*θ*)^⊤^ = −*Q*(*θ*). *If a cross-sectional assay draws one observation per cell at stationarity, then*

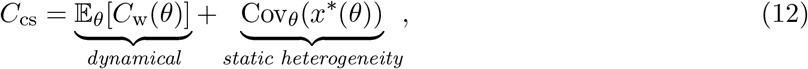

*and both terms are positive semidefinite*.

*Proof*. Immediate from the law of total covariance, Cov(*x*) = E[Cov(*x* | *θ*)] + Cov(E[*x* | *θ*]), with Cov(*x* | *θ*) = *C*_w_(*θ*) and E[*x* | *θ*] = *x*^∗^(*θ*). □

Equation (12) is the intrinsic–extrinsic decomposition [46, 51] written in continuous form. The first term is what the Lyapunov inversion requires; the second is frozen parameter variation carrying no dynamical information at all. Dual-reporter assays exist precisely to estimate the split [47, 48].

#### Corollary 4.2

(Direction of the bias). *Suppose C*_cs_ *is substituted for C in the Lyapunov equation with the correct Q. Then in the scalar case the inferred relaxation rate satisfies*

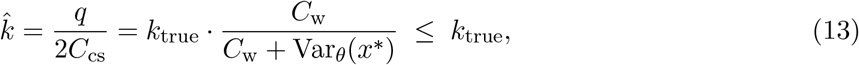

*with equality only if the population is homogeneous. Inferred relaxation is therefore always too slow, and the system appears closer to marginal stability than it is*.

#### Why the direction matters

A downward bias in |Re *λ*| is a downward bias in the estimated distance to bifurcation. Cross-sectional data therefore make systems look closer to a critical transition than they are. Where such estimates inform a decision about whether an intervention window remains open, the error is in the unsafe direction, and it is not small: heterogeneity comparable to within-cell variability (routine in single-cell data) already inflates the apparent relaxation time by a factor of two to three.

### 4.4 Genuine Non-Ergodicity: Slow and Glassy Modes

Condition E2 fails outright for a class of biological processes. Epigenetic marks (DNA methylation, histone modifications, chromatin state) evolve on timescales far longer than transcriptional or signalling dynamics and are heritable across divisions, so the state space is not explored within the lifetime of the experiment or the organism.

The phenomenology is that of glassy physics [71, 72]: a rugged landscape with many metastable minima, waiting times drawn from heavy-tailed distributions, subdiffusive spreading ⟨Δ*E*(*t*)^2^⟩ ∼ *t*^*α*^ with *α <* 1, power-law rather than exponential memory kernels, and ageing — a response that depends on the system’s history rather than only its current state. The last of these is a violation of the fluctuation–dissipation relationship, which removes the very principle that licenses passive fluctuation measurement as a proxy for response.

Empirically, the variance of methylation marks grows with age rather than saturating, which is the basis of epigenetic age estimators [73, 74], and single-cell methylation data show increasing heterogeneity with age and in senescence. In such regimes there is no stationary distribution to estimate: the time average does not converge, and the ensemble average measures a distribution over frozen states rather than a dynamical covariance.

#### Position taken here.

Modes whose ergodic time exceeds the observation window are declared outside the operator’s validity domain and are reported as such. They are not estimated with wide error bars; they are excluded, because the quantity they would estimate does not exist. Section 14 makes this declaration an explicit term of the exported contract.

### 4.5 A Three-Regime Taxonomy

Table 6 summarises the resulting decision procedure.

### 4.6 Diagnostics: Testing Rather Than Assuming

Ergodicity is testable, and should be tested.

#### Ergodicity-breaking parameter

For a time-averaged observable *δ*^2^(*T* ) computed over windows of length *T*, define

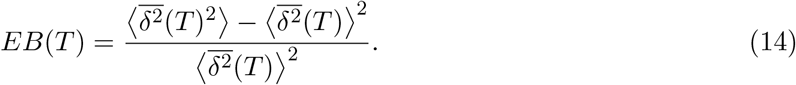

For an ergodic process *EB*(*T* ) → 0 as *T* → ∞; for a weakly non-ergodic process it approaches a nonzero constant. This statistic is standard in single-particle tracking [75–77] and transfers directly.

#### Time-averaged versus ensemble-averaged mean-squared displacement

Equality of the two, for the same lag, is the operational definition of ergodicity; systematic divergence with increasing measurement time signals ageing [76].

#### Variance growth

If Var(*x*(*t*)) grows with *t* rather than saturating, no stationary distribution exists and the operator does not apply.

#### Direct comparison

Where a small longitudinal panel accompanies a large cross-sectional dataset, the difference *Ĉ*_cs_ −*Ĉ*_long_ estimates Cov_*θ*_(*x*^∗^) directly, calibrating the correction in Theorem 4.1 without requiring longitudinal data at scale.

#### Dual-reporter designs

Two identical reporters driven by identical promoters partition variability: the covariance between them estimates the shared (extrinsic and static) component, and half the mean squared difference estimates the intrinsic component [46, 47]. Where such constructs are available they measure the decomposition of Theorem 4.1 directly. Allele-resolved single-cell measurements serve the same purpose in natural systems.

### 4.7 Recovering Dynamical Information from Snapshots

The most consequential limitation of cross-sectional data is the absence of lag information. Three routes partially recover it.

#### Velocity statistics

Metabolic labelling and splicing-based approaches now furnish, for each cell in a snapshot, an estimate of the *rate of change* as well as the state [78–81], and this information has been used to estimate Jacobians from single-cell data [82]. The relevant identity is

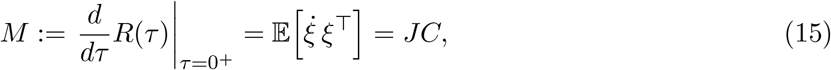

which combined with 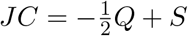 (Theorem 5.2) gives both unknowns at once:

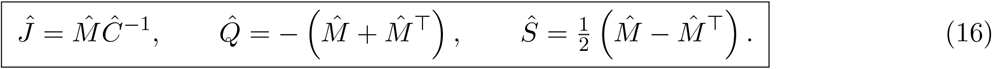

A single snapshot bearing per-cell velocities identifies the generator, the diffusion matrix, *and* the probability current, provided the ensemble is homogeneous in the sense of E3. We verified (16) numerically to machine precision on randomly generated systems with strongly anisotropic *Q*.

#### Synchronised perturbation ensembles

Applying a perturbation to a synchronised population and sampling the ensemble at successive times yields the response function directly, from which eigen-values follow by exponential fitting (Section 12.4). This substitutes an active for a passive measurement and sidesteps the fluctuation–response assumption entirely.

#### Pseudotemporal ordering

Trajectory-inference methods order cells along a putative progression [83, 84]. We note this route but do not rely on it: pseudotime is identified up to a monotone reparameterisation, so it does not deliver the absolute rates that eigenvalues represent, and the ordering itself presupposes that the population traverses a common path, a strong form of E3.

##### Remark 4.3

(Ergodicity is prior to everything else). The velocity route removes the need for time series but not the need for E3. A snapshot of a heterogeneous population yields *Ĉ* inflated by Cov_*θ*_(*x*^∗^) and 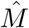 contaminated by systematic differences in *x*^∗^(*θ*), and no amount of velocity information repairs this. Stratification or explicit mixture modelling must precede the inversion, not follow it.

## 5 Lyapunov Equation and the Structure of the Inverse Problem

### 5.1 Derivation

At stationarity 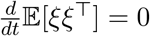, which with (3) yields the continuous Lyapunov equation

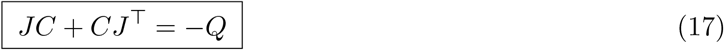

with explicit solution 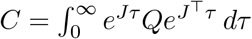, unique and positive definite when *J* is Hurwitz and (*J, B*) is controllable. Equation (17) states a balance: deterministic contraction offsets stochastic injection, and what remains is the observable covariance.

#### Remark 5.1

(Rank-deficient *Q*). By Proposition 3.1(ii), conservation laws make *Q* singular. Then *C* ≻ 0 requires (*J, B*) controllable, i.e. every direction must be reachable by noise propagated through the dynamics even if not directly excited. Where this fails, *C* is singular, *C*^−1^ does not exist, and both Theorem 5.2 and (16) must be restricted to the reachable subspace. Consequently, this approach accurately manages conserved total pools by eliminating them prior to matrix inversion rather than retaining them throughout the calculation.

### 5.2 The Inverse Problem is Underdetermined

Equation (17) relates symmetric matrices and so supplies *n*(*n* + 1)*/*2 scalar constraints for the *n*^2^ unknowns of *J*. The deficit is *n*(*n*−1)*/*2. Counting establishes that a problem is underdetermined but not what the undetermined directions are; here they are exactly characterisable, and the characterisation is severe enough that stating it is the central technical step of this article.

#### Theorem 5.2

(Gauge freedom of the Lyapunov inverse). *Let C* ≻ 0 *and Q* = *Q*^⊤^ *be given. Then J satisfies JC* + *CJ*^⊤^ = −*Q if and only if*

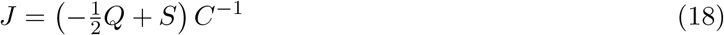

*for some antisymmetric S* = −*S*^⊤^. *The solution set is an affine family of dimension n*(*n* − 1)*/*2. *Proof*. (⇐) Substituting (18) gives 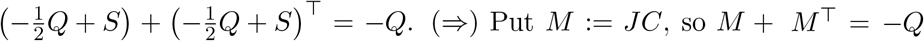. Define 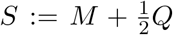; then *S* + *S*^⊤^ = *M* + *M*^⊤^ + *Q* = 0, so *S* is antisymmetric and 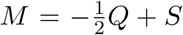. Since *C* is invertible, *J* = *MC*^−1^. The space of real antisymmetric *n* × *n* matrices has dimension *n*(*n* − 1)*/*2.

#### Corollary 5.3

(The spectrum is not identifiable from (*C, Q*)). *The map* 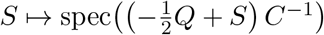 *is generically non-constant. The eigenvalues of J (including the number of oscillatory modes, their frequencies and their damping ratios) are therefore* not *determined by the stationary and noise covariances*.

This contradicts a claim made frequently in literature. It is not the case that *J* is unidentifiable while its spectrum is identifiable; the spectrum is unidentifiable too.

### 5.3 The Gauge is the Probability Current

The antisymmetric matrix *S* is not an artefact. By (15) it is the antisymmetric part of 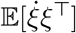, which is the stationary probability current: the circulating flux in state space. When *S* = 0 the process is reversible and detailed balance holds. When *S* ≠ 0 the system sustains cycles.

Biological systems sustain cycles by construction. Maintaining organisation requires continuous free-energy dissipation, so cellular networks operate in non-equilibrium steady states with non-zero entropy production [85, 86], and the trade-off between precision and dissipation is quantified by the thermodynamic uncertainty relation [87]. Phosphorylation–dephosphorylation cycles, transcription– degradation cycles, and molecular motors all carry net flux. Broken detailed balance has been measured directly in cellular systems [88] and reviewed extensively [89, 90].

#### Consequence.

*S* ≠ 0 is the generic biological case, not an edge case. The gauge freedom of Theorem 5.2 is therefore not a technicality to be waved away by a genericity assumption. It is the normal situation, and it must be resolved with data rather than with an assumption.

#### Remark 5.4

(The optical reading). Theorem 5.2 admits the interpretation anticipated in Section 1.8. By (1) the stationary covariance is the frequency-integrated power 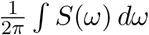, so an inversion based on *C* alone attempts to reconstruct a source’s line structure from its total brightness. That this fails is not surprising; what Theorem 5.2 adds is the exact dimension of the failure, *n*(*n* − 1)*/*2, and the identification of the lost coordinate as the probability current. Restoring *τ* - or *ω*-resolution (Routes 2 and 3 of Section 5.4) is the methodological equivalent of dispersing the beam, and it recovers everything that was lost.

### 5.4 What Restores Identifiability

Four routes exist. The operator of Section 6 uses the second and third. The first is recorded mainly to show why it is unsuitable.

**Route 1: assume reversibility**. Setting *S* = 0 gives 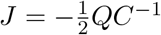, uniquely determined.

#### Proposition 5.5.

*If S* = 0 *and Q* ≻ 0 *then* 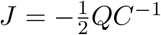 *is similar to a symmetric negative-definite matrix, so its spectrum is real and strictly negative*.

*Proof*. 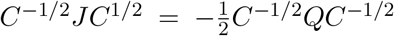, symmetric and negative definite; similarity preserves the spectrum.

Assuming detailed balance thus *forbids oscillation*. Since endocrine axes, cell cycles and circadian circuits oscillate, this route is unsuitable as a default for biology, though it remains reasonable for near-equilibrium subsystems such as rapid binding equilibria.

**Route 2: lagged covariance**. The autocovariance

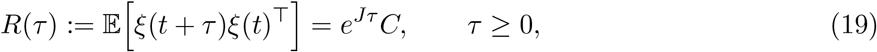

carries what *C* lacks.

#### Proposition 5.6

(Identifiability from a single lag). *Let C* ≻ 0 *and let τ >* 0 *satisf*(*y τ* |Im(*λ*_*i*_ )| *< π for all eigenvalues of J. Then J is uniquely determined by* (*C, R*(*τ* )) *via J* = *τ*^−1^Log (*R*(*τ* )*C*^−1^ ), *with* Log *the principal matrix logarithm*.

*Proof*. From (19), *e*^*Jτ*^ = *R*(*τ* )*C*^−1^. The frequency condition ensures no eigenvalue of *e*^*Jτ*^ lies on the closed negative real axis, so the principal logarithm is well defined and inverts the exponential on the relevant branch.

In practice the logarithm is not applied directly, since it amplifies estimation error; the lag information enters as a fitting term, as in (22).

**Route 3: velocity statistics**. Equation (16). This is the cross-sectional analogue of Route 2 and is the appropriate choice when per-cell rate estimates are available.

**Route 4: impose structure**. Constraining *J* to a known sparsity pattern reduces the unknown count from *n*^2^ to the number of admissible edges. If that falls below *n*(*n*+1)*/*2 the system may become identifiable from equal-time data alone. This is the route taken in the metabolomics literature, using stoichiometric reconstructions to fix the support of *J* [26, 28, 29], and is developed in Section 10.5. Its identifiability is conditional on the assumed topology being correct.

### 5.5 Joint Recovery of the Diffusion Matrix

Routes 2 and 3 have a consequence that materially changes the practical status of the noise model. Once *J* is identified, *Q* follows:

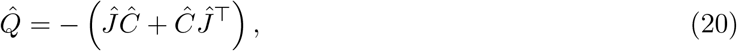

or equivalently, from velocity data, 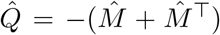. The diffusion matrix therefore need not be assumed, hard-coded, or transferred between datasets. It is estimated, per dataset, from the same measurements that determine the generator, which is the correct response to the observation that noise structure is dataset-dependent (Section 3.4). This yields a falsification test at no additional cost.

**Protocol 5.7** (Positive-semidefiniteness test). Compute 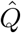 by (20) and examine its spectrum. If 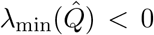 beyond estimation error, the model (3) is misspecified. The leading candidate explanations, in order of prior plausibility for biological data, are: (i) the driving noise is coloured rather than white (typically extrinsic modulation, requiring the augmentation (7)); (ii) unobserved variables couple to the measured ones; (iii) the process is non-stationary over the estimation window; (iv) the linear approximation has failed.

**Protocol 5.8** (White versus coloured noise). For white driving noise, *R*(*τ* ) = *e*^*Jτ*^ *C* has a one-sided derivative *JC* ≠ 0 at the origin, so the autocovariance has a corner at *τ* = 0. For noise with correlation time *τ*_*c*_ *>* 0, *R*(*τ* ) is differentiable at the origin with *R*^*′*^(0) = 0, so the autocovariance is smooth there. Examining the behaviour of 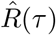 as *τ* → 0 therefore discriminates the two cases and estimates *τ*_*c*_, which sets the required augmentation in (7).

#### Summary of the identifiability position.

Equal-time covariance alone determines *J* only under detailed balance, which forbids oscillation and is biologically atypical. UNS is therefore defined on (*C, R*(*τ* )) or on (*C, M* ), optionally augmented by structural constraints. In either case *Q* is recovered rather than assumed, and its positive semidefiniteness is a model-adequacy test. Where only equal-time data exist, the operator must report gauge-invariant quantities and declare the remainder unidentified rather than returning an arbitrary member of the family (18).

## 6 The UNS Operator

### 6.1 Definition

#### Definition 6.1

(UNS operator). Let ℋ_*A*_ denote the set of Hurwitz matrices whose sparsity pattern is contained in an admissible support *A* (with *A* = **1** imposing no structural constraint). Given a stationary covariance estimate *Ĉ*, dynamical information in the form of lagged covariances 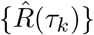 or a velocity cross-moment 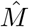, the UNS operator is

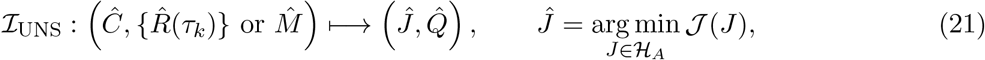

with

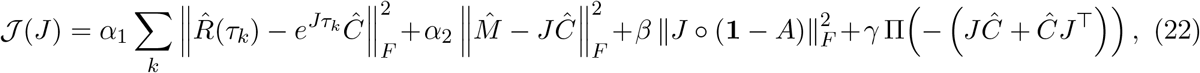

where Π(·) penalises departure from positive semidefiniteness of the implied diffusion matrix, and 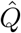 is then recovered by (20). Weights are set by cross-validation on held-out segments or cells.

Two features distinguish (22) from a naive Lyapunov inversion. The stationarity residual 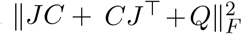 does not appear as a separate term, because with *Q* recovered by (20) it is identically zero and carries no information. Its role is taken by the semidefiniteness penalty, which is the part of the stationarity constraint that retains content. And the dynamical terms are not optional refinements: by Corollary 5.3, setting *α*_1_ = *α*_2_ = 0 returns the problem to the degenerate family of Theorem 5.2.

A third feature is worth stating because it is not apparent from the form: (22) never inverts *Ĉ*. Both dynamical residuals multiply by *Ĉ* rather than by *Ĉ*^−1^, so the objective remains well defined when the covariance estimate is singular (the usual case, for the reasons set out in Section 7.3). The closed-form routes of Proposition 5.6 and (16) do not share this property.

### 6.2 Setting the Weights, and the Two Terms That Carry Assumptions

Cross-validation on held-out segments or cells fixes the four weights, but two of the terms carry inputs that must themselves be estimated or asserted, and cross-validation will not detect a bad choice of either: it selects the weight that best predicts held-out data *given* the input, and a systematically wrong 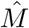 or a topology-shaped *A* will simply be assigned a weight that minimises the damage.

#### The velocity term: 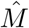 must be reported as an estimate

The identity 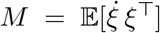 is exact, the estimator is not. Where 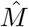 comes from a perturbation design rather than from percell velocities, its form matters. Two natural constructions (a symmetrised difference estimator and a Gram-type estimator formed from the perturbation and response matrices) differ in bias and in how they degrade as perturbation coverage falls, and the ordering between them reverses: on our own factorial comparison the symmetrised form is preferable at high coverage, while below roughly 70% coverage the Gram form is more accurate, and doubling the replicate budget buys about three times what the choice of form buys. Neither the estimator nor the coverage is a free parameter of the analysis, and both belong in D.

#### The support term: *A* is a hypothesis, and it must be tested as one

The penalty 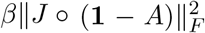 is Route 4 of Section 5.4, whose identifiability is conditional on the assumed topology being correct. Where *A* is taken from a curated interaction database that condition is a substantive empirical claim, and two failure modes are common enough to warrant explicit checks.

##### A support prior must be tested against a structure-preserving null, not against no prior at all

Curated networks have strongly heterogeneous degree, and a degree sequence alone induces covariance structure. The relevant comparison is therefore against an ensemble that preserves the degree sequence and randomises the edges (degree-preserving rewiring) and a support prior that does not beat that ensemble has contributed its degree distribution and nothing else, however large its improvement over *β* = 0. This is a distinct test from cross-validating *β*, and passing the second does not imply passing the first.

##### Improvement must be placed on the interval the design admits

A support prior that improves held-out fit has demonstrated only that the improvement is non-zero. What matters is where it sits between the null and the design-matched ceiling of Remark 14.1: a prior recovering a small fraction of the attainable range is not evidence that the operator works, and a narrow interval inflates the ratio, so the absolute improvement must be reported alongside it.

We record the expectation this sets up, since it is testable and we regard it as the sharpest available check on the operator. If the equal-time corner *α*_1_ = *α*_2_ = 0 attains only a small fraction of the design-matched range under the best available *A* (as we find for a human proteomic dataset in work reported separately) then the dynamical terms are not refining a mostly-correct inference but supplying most of it, and a study adding lagged or velocity data should show a correspondingly large movement in *π*. A design that adds such data and does *not* move *π* has falsified something, and the operator is the leading candidate.

### 6.3 Reduction to an Invariant Subspace

#### Definition 6.2

(Abstraction operator). Order the eigenvalues of *Ĵ* by increasing |Re(*λ*_*i*_)| and let *r* be the number retained, chosen so that no conjugate pair is split. Let

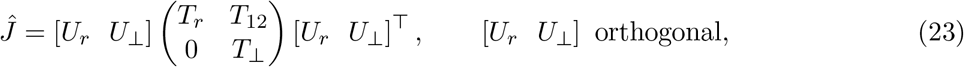

be a real Schur decomposition ordered so that spec(*T*_*r*_) is the retained set; *T*_*r*_ is real quasi-triangular, with complex modes in conjugate 2 × 2 blocks. Then *ĴU*_*r*_ = *U*_*r*_*T*_*r*_, so col(*U*_*r*_) is an invariant subspace, and the abstraction operator is

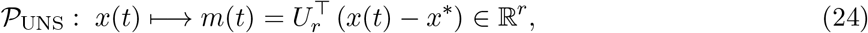

with reduced generator, projected noise and reconstruction

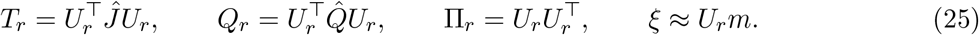

The projection and the reconstruction use the same factor, because *U*_*r*_ is orthonormal. This is the practical advantage of (23) over the eigenvector form of Remark 6.3: it is computed by a backward stable algorithm, it does not require the retained eigenvalues to be simple or even distinct, and it degrades continuously as two of them approach each other.

#### Remark 6.3

(The eigen-refinement, and the dictionary between the two forms). Where spec(*T*_*r*_) is simple and well separated relative to Σ_*T*_, the reduced generator may be diagonalised, *T*_*r*_ = *Y* Λ_*r*_*Y* ^−1^, giving right and left modal bases

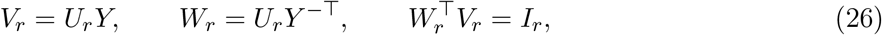

in which 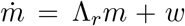 is diagonal to linear order. This is the form in which the description is *interpreted* : *v*_*i*_ identifies which variables participate in mode *i, w*_*i*_ which must be perturbed to excite it, and the distinction governs actuator placement (Section 14).

Two facts about the relationship should be stated, because they are easy to get wrong in either direction.

*The two projectors are different operators*. 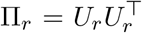 is the orthogonal projector onto col(*U*_*r*_); the spectral projector of the full generator, 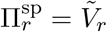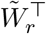 built from the right and left eigenvectors of *Ĵ* itself, projects onto the same subspace but *along the discarded invariant subspace*, which for non-normal *Ĵ* is not the orthogonal complement. Consequently 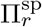 commutes with *Ĵ* and decouples the dynamics exactly, while Π_*r*_ does not and leaves the residual coupling *T*_12_ of Remark 6.4. The trade is stability against exactness, and it is resolved here in favour of stability, because 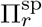 is undefined when the retained and discarded spectra touch, the case a defective or near-defective generator produces, and precisely the case Section 3.6 argues is not exotic in biology.

#### Non-normality is not lost by the change of form; it moves

Since *U*_*r*_ is orthonormal, ∥Π_*r*_∥_2_ = 1 identically, so the projector norm is no longer an informative diagnostic and is replaced by the two quantities of Remark 6.6.

For normal *Ĵ* the two forms coincide up to a rotation: *T*_*r*_ is then block diagonal, *Y* orthogonal, *V*_*r*_ = *W*_*r*_, and 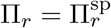. The distinction is drawn here because Section 3.6 establishes that biological generators are not normal.

##### Remark 6.4

(The descriptor is a restriction, not a closed reduced model). Invariance of col(*U*_*r*_) gives the zero block below the diagonal in (23) but not *T*_12_ = 0. Writing *x* − *x*^∗^ = *U*_*r*_*m* + *U*_⊥_*z*,

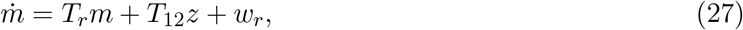

so the retained coordinates are driven by the discarded ones through *T*_12_. The marginal process in *m* has the law of the autonomous Ornstein–Uhlenbeck process 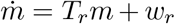 under conditions of which *T*_12_ = 0 and Cov(*m, z*) = 0 are two structurally distinct sufficient cases; neither is necessary, and they should not be conflated. Away from them the marginal is non-Markovian.

The consequence for reporting is that (*T*_*r*_, *U*_*r*_, *Q*_*r*_) is the restriction of 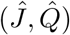 to an invariant sub-space, not a closed model of a subsystem. Supplying closure requires a realisation of the discarded block, not a summary of it, and ∥*T*_12_∥ belongs in the exported qualification for exactly this reason. The same caveat applies verbatim to the eigen-refinement, where the coupling reappears as the off-diagonal block of *Ĵ* in the biorthogonal basis.

##### Remark 6.5

(Basis freedom, and what to compare across studies). For orthogonal *O* ∈ ℝ ^*r×r*^ the triple (*T*_*r*_, *U*_*r*_, *Q*_*r*_) → (*O*^⊤^*T*_*r*_*O, U*_*r*_*O, O*^⊤^*Q*_*r*_*O*) describes the same dynamics, so descriptors from different studies are comparable only modulo this equivalence unless an ordering convention is fixed, and no convention stated in terms of *T*_*r*_ alone is canonical when *T*_*r*_ has normal 2 × 2 blocks, ties in |Re *λ*|, or repeated eigenvalues. The basis-independent quantities should therefore always be reported alongside: spec(*T*_*r*_), principal angles between col(*U*_*r*_) and a comparator subspace, ∥*T*_12_∥, and Tr *Q*_*r*_. In the eigen-refinement the corresponding freedom is *Y* → *Y T* for *T* a permutation and a diagonal scaling, under which Λ_*r*_ is fixed and *Q*_*r*_ ↦ *T*^−1^*Q*_*r*_*T*^−⊤^.

##### Remark 6.6

(Two basis-independent measures of non-normality). Under (23) the projector is orthogonal and ∥Π_*r*_∥_2_ = 1 identically, so non-normality must be reported by quantities read off *T* itself. Two suffice, and both are invariant under orthogonal similarity.

#### Departure from normality

Henrici’s measure 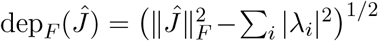 equals the Frobenius norm of the strictly upper triangular part of the complex Schur form, is zero exactly for normal *Ĵ*, and is what bounds the gap between spectral and pseudospectral behaviour [64].

#### Coupling into the retained block

∥*T*_12_∥_*F*_ measures how strongly the discarded coordinates drive the retained ones (Remark 6.4). It is the quantity that decides whether the descriptor may be read as a subsystem model, and it has no analogue in the projector norm.

On a randomly generated symmetric generator both vanish: dep_*F*_ = 0.000, ∥*T*_12_∥_*F*_ = 0.000. On a non-normal generator of the same dimension, dep_*F*_ = 3.21 against ∥*Ĵ*∥_*F*_ = 8.71, with ∥*T*_12_∥_*F*_ = 2.98, a coupling of the same order as the reduced generator itself. Where the eigen-refinement of Remark 6.3 is also reported, *κ*(*Y* ) bounds by (11) how far the retained eigenvalues can move under model error, and the oblique projector norm 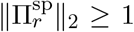 measures the angle between the retained and discarded invariant subspaces; on the two generators above these are 1.000 and 1.000 for the symmetric case and, on a further non-normal draw, 9.74 and 2.61 respectively.

##### Remark 6.7

(Why not a rank constraint). Defining the reduction as a minimisation over matrices of rank at most *r* is not well posed. A matrix of rank *r < n* has *n* − *r* zero eigenvalues. The associated process has non-decaying directions and admits no stationary covariance, so the residual is bounded away from zero along the kernel and the minimiser is not interpretable as a generator. Moreover 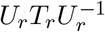 is undefined, since *U*_*r*_ ∈ ℝ^*n×r*^ is not invertible. The correct object is an *r*-dimensional invariant subspace with the orthogonal projection (25).

##### Remark 6.8

(Mode selection). According to Section 3.5, modes are ordered by |Re(*λ*_*i*_)| obtained from *Ĵ*, never by the magnitude of the corresponding covariance eigenvalue. Retention additionally requires that the mode be resolvable: its eigenvalue must be separable from the estimation noise floor under the resampling scheme of Section 7, and its ergodic time must satisfy regime E of Table 6.

### 6.4 Exported Quantities

#### Dimension

The state dimension falls from *n* to *r*, typically from 10^2^–10^3^ to 2–5.

#### Timescale preservation

spec(*T*_*r*_) is exactly the retained subset of the eigenvalues of the full generator, and is unchanged by the basis freedom of Remark 6.5.

#### Decoupling, conditionally

Where the eigen-refinement of Remark 6.3 applies, the dynamics are diagonal in modal coordinates to linear order. Where it does not (coalescing or clustered eigenvalues) *T*_*r*_ remains block quasitriangular and the description is of a subspace rather than of separated modes.

#### Interpretability

Under the eigen-refinement, *v*_*i*_ identifies which variables participate in mode *i* and *w*_*i*_ which must be perturbed to excite it; the distinction governs intervention (actuator) placement. Without it, col(*U*_*r*_) still identifies the combinations of variables the description is about.

#### Noise calibration

*Q*_*r*_ is inherited from the *recovered* 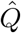, so modal fluctuation amplitudes are correct without a noise assumption.

#### Restriction, not closure

∥*T*_12_∥_*F*_ is reported, because it decides whether the exported object may be read as a subsystem model at all (Remark 6.4).

#### Conditioning

dep_*F*_ (*Ĵ*) is reported for departure from normality. Where the eigen-refinement is used, *κ*(*Y* ) is reported alongside it and bounds by (11) how far eigenvalues can move under model error (Remark 6.6).

## 7 Identifiability, Sample Complexity, and Uncertainty

### 7.1 Four Levels of Identifiability

Four distinct questions are frequently conflated, and a quantity must clear all four before it may be exported.

#### Ergodic admissibility

Does the quantity refer to a stationary object that the available data can estimate at all? Section 4.

#### Gauge identifiability

Is the quantity invariant across the family (18)? Section 5.2.

#### Structural identifiability

Given noiseless data of the assumed type in unlimited quantity, is the parameter unique? Analysable by differential algebra [91–93].

#### Practical identifiability

Given the finite noisy data actually available, can it be estimated usefully? Profile-likelihood analysis is standard [94, 95]. For stochastic kinetics specifically, the linear noise approximation supports a Fisher-information treatment [21, 96].

### 7.2 Sample Complexity for Both Data Types

Feasibility is governed by how much data are required, and the two data types have different requirements.

#### Longitudinal

Observations separated by less than the correlation time are not independent. For a record of duration *T* sampled finely, the effective sample size for the slowest retained mode is

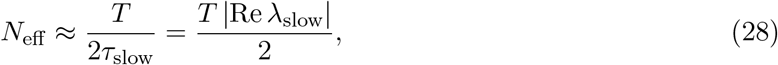

so resolving an *n*-dimensional covariance requires *T* ≳ 2*nτ*_slow_. Sampling faster does not help; only observing longer does. If the slow mode of a circuit has a timescale of weeks, even a ten-dimensional description requires months of monitoring.

#### Cross-sectional

Cells are independent, so *N*_eff_ = *N*_cells_ and the covariance is easy to estimate, which is precisely why the cross-sectional route is attractive, and why the bias of Theorem 4.1 is dangerous: it is a bias, not a variance, and it does *not* shrink as more cells are sequenced. Increasing *N* produces an increasingly precise estimate of the wrong quantity.

##### The asymmetry that matters.

Longitudinal data are variance-limited: more observation time buys accuracy. Cross-sectional data are bias-limited: more cells buy precision around a displaced target. No amount of sequencing depth substitutes for stratification or for the velocity information of Section 4.7.

### 7.3 The Rank-Deficient Regime, and Why it is the Usual One

Every result in Section 5 assumes *Ĉ* ≻ 0: Theorem 5.2 inverts *C*, Proposition 5.6 forms 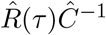, and the velocity recovery (16) forms 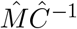. For a sample covariance built from *N* observations of *n* variables, rank *Ĉ* ≤ min(*N* − 1, *n*), so the hypothesis fails outright whenever *N* ≤ *n*, which is the standard shape of the assays the operator is aimed at. A transcriptomic panel of 10 –10 genes across a few hundred conditions, or a proteomic covariance over 10^3^ proteins across a few hundred cell lines, is not a marginal case: it is the case.

The hypothesis is therefore not a technical convenience and cannot be left implicit. Two distinct kinds of rank deficiency arise, they call for opposite numerical responses, and conflating them accounts for a class of published null results.

#### Structural deficiency

*C* is exactly singular by construction, with a kernel that is known. This occurs when conservation laws leave ℓ^⊤^*S* = 0 (Proposition 3.1(ii)) and when a diffusive generator is built on a graph Laplacian, whose kernel has dimension equal to the number of connected components. Here the Moore–Penrose pseudo-inverse is not a numerical expedient but the correct object: the dynamics live on the complement of the kernel, and *C*^+^ is the exact inverse there. Applying a dense solver instead returns the kernel amplified by rounding error, and the result is indistinguishable from noise. In the demonstration below, recovering a graph from *C* = *L*^+^ scores 1.000 by AUROC through the pseudo-inverse and 0.516 through the solver.

#### Sampling deficiency

*C* is generically full rank in the population but its estimate is not, because *N* ≤ *n*. Here the pseudo-inverse is *not* the fix, and this is the point most easily missed: *Ĉ*^+^ inverts the *N* − 1 sample directions and annihilates the remaining *n* − *N* + 1, so it is a well-conditioned estimate of the wrong operator. It removes the numerical blow-up while leaving (and in the table below worsening) the statistical failure. The required response is regularisation: a ridge or shrinkage estimator, or a penalised inverse where sparsity of the precision matrix can itself be justified, with the regularisation weight chosen on held-out data and reported as part of the estimation protocol rather than absorbed silently.

##### A null result that is an artefact.

At *N/n* ≤ 1 the sample covariance is exactly singular, yet a dense solver need not fail: it returns entries of order 1*/*(*ε*_mach_∥*Ĉ*∥) (10^15^–10^18^ at unit noise scale in our runs) and an AUROC of ≈ 0.50, which is a clean, publishable-looking statement that the network carries no information about the covariance. The magnitude is set by machine precision and the units of the data, not by the network. The same data through a shrinkage estimator score 0.78–0.90. Any negative result about fluctuation-based network inference must therefore report rank *Ĉ* (with the tolerance used), *N/n*, the reciprocal condition estimate of *Ĉ*, and the inverse actually used, or it cannot be distinguished from this failure.

#### Consequences for the operator

Three of them, and they should be stated in *D*.

First, Definition 6.1 is well posed without modification (it never forms *Ĉ*^−1^), since the lag residual uses 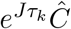 and the velocity residual *JĈ*, both of which multiply rather than invert. This is a substantive advantage of the variational form over the closed-form routes and was not the reason for adopting it. The closed forms (16) and Proposition 5.6, by contrast, require the regularised inverse and inherit its bias.

Second, in the sampling-deficient regime the gauge freedom of Theorem 5.2 is joined by a second and larger indeterminacy: *Ĵ* is unconstrained on the *n* − *N* + 1 directions the sample does not span, whatever dynamical information is supplied. The honest report is the intersection (the identified subspace is at most (*N* − 1)-dimensional) and *r* must be chosen well inside it.

Third, the sample-complexity asymmetry of the previous subsection acquires a floor. For cross-sectional data *N*_eff_ = *N*_cells_ is large and this constraint is usually slack; for longitudinal data *N*_eff_ ≈ *T/*(2*τ*_slow_) by (28), so rank *Ĉ* is bounded by the observation duration in units of the slowest retained timescale, and no sampling rate repairs it. Resolving an *n*-dimensional covariance requires *T* ≳ 2*nτ*_slow_ for the same reason it requires *N* ≳ 2*n* cross-sectionally, and Table 8 indicates that the constant is not far from 2.

**Table 8:** Recovery of a 200-node Erdős–Rényi graph (Pr[edge] = 0.03) from the precision matrix, under isotropic forcing, as a function of the observation budget. AUROC over all non-diagonal pairs; 0.5 is chance. *Ridge* is (*Ĉ*+*ε n*^−1^ Tr *Ĉ I*)^−1^ with *ε* = 10^−2^; *LW* is the Ledoit–Wolf shrinkage estimator. Read across the *N/n* ≤ 1 rows: the dense solver returns chance, the pseudo-inverse returns chance or worse, and only regularisation recovers usable structure. Read down: the identifiability the population case promises requires *N/n* ≳ 2 before any estimator delivers it.

| $N$ | $N/n$ | $\text{rank } \hat{C}$ | $\hat{C}^{-1}$ | $\hat{C}^{+}$ | ridge | LW |
| --- | --- | --- | --- | --- | --- | --- |
| population | — | 200 | 1.000 | 1.000 | — | — |
| 60 | 0.30 | 59 | 0.500 | 0.373 | 0.754 | 0.780 |
| 150 | 0.75 | 149 | 0.488 | 0.635 | 0.752 | 0.877 |
| 200 | 1.00 | 199 | 0.535 | 0.571 | 0.745 | 0.904 |
| 400 | 2.00 | 200 | 0.919 | 0.919 | 0.925 | 0.970 |
| 1000 | 5.00 | 200 | 0.995 | 0.995 | 0.996 | 0.997 |
| 4000 | 20.00 | 200 | 1.000 | 1.000 | 1.000 | 1.000 |

### 7.4 Uncertainty Quantification

Three mechanisms are used, and their outputs form part of the exported description. *Block bootstrap*: resample longitudinal records in blocks longer than the correlation time, or resample cells with stratification, recompute the inputs, and re-run I_UNS_. *Bayesian posterior* : place a prior on *J* informed by topology as in (36) and sample the posterior implied by (22). *Fisher information*: the Cramér–Rao bound gives asymptotic variance floors and, through the eigenstructure of *I*(*θ*), identifies the sloppy directions along which estimation is hopeless [4, 96].

Uncertainty is not decoration. A mode with *λ* = −0.1 ± 0.05 and one with *λ* = −0.1 ± 0.5 have entirely different control implications: the first admits an aggressive controller, the second demands robust design against a process whose stability is in question.

## 8 Local Linearizability as a Structural Property

### 8.1 Homeostasis Enforces the Operating Regime

Local linearization is usually described as an assumption, and for generic nonlinear systems that description is apt. For physiological circuits the situation differs in a way that strengthens the case for the operator.

Physiological circuits are regulated by feedback whose explicit function is to hold operation near a setpoint [97, 98]. Blood glucose is held within a narrow band despite large meal perturbations; core temperature varies by less than one percent of its absolute value; plasma osmolality is regulated within a few percent. The condition required for linearization (small excursions about a fixed point) is not an approximation the modeller imposes but a state the organism actively maintains.

The formal theory has developed considerably. Singularity-theoretic treatments classify the network structures capable of holding an output invariant under input variation, and identify when the invariance is structural rather than parametric [99–101]. Complementary work characterises the topologies achieving robust adaptation [102] and the biomolecular implementations realising integral feedback [103]. Linearizable operation is a designed property of regulatory architecture, not a modelling convenience.

### 8.3 Dynamical Compensation

The same feedback induces *dynamical compensation*: internal mechanisms adjust to preserve macroscopic outputs despite parameter drift [104, 105]. In a hormone axis where secretion rate *s* and clearance rate *d* both vary, feedback that adjusts *s* to track *d* preserves *x*^∗^ = *s/d* under proportional variation, and an observer measuring only *x*^∗^ cannot distinguish (*s, d*) from (2*s*, 2*d*).

Compensation suppresses sensitivity along specific parameter directions, rendering parts of the linearized dynamics weakly observable. The Jacobian exists and is well defined; certain combinations of its entries are simply absent from the data. The appropriate response is to report this rather than conceal it: the operator returns a reduced description together with the statement that uncertainty is large orthogonal to the identified directions.

### 8.3 The Evolutionary Origin of Low Rank

The premise that biological covariance is dominated by a small number of modes is usually offered as an empirical observation. It has a stronger foundation. Fluctuation–response analyses show that evolution under selection tends to produce a *dominant mode*: a single direction along which both spontaneous fluctuation and response to environmental change are concentrated [106–108]. Selection for robustness in most directions, combined with plasticity along the direction of adaptation, generates the spectral anisotropy that (9) then amplifies.

If this account is correct, the low-rank structure the operator exploits is a signature of evolutionary history rather than a convenient approximation. Note that it also supplies an independent reason to expect 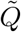 to be anisotropic: the noise and the dynamics are shaped by the same selective process and are not statistically independent.

### 8.4 Multistability

Physiological systems frequently exhibit multistability. As the system approaches a basin boundary, the dominant eigenvalue converges toward zero, relaxation times experience critical slowing down, and any single linearization ceases to be globally valid. Consequently, the operator does not assume *a priori* the existence of a viable linear model; instead, it dynamically determines which components of the local linear structure are supported by observed fluctuations, establishing both their temporal window of validity and their associated uncertainty.

## 9 Topological Constraints on Inferred Spectra

Network topology imposes fundamental constraints on inferred eigenvalues, thereby providing empirical falsification tests that are inaccessible through purely statistical inference. Because biological networks are inherently directed, signed, and non-normal, several analytical bounds conventionally applied in this context are rendered invalid. Consequently, this section employs mathematical frameworks robust to these topological complexities.

### 9.1 Preliminaries

Represent the network as a graph with weighted adjacency matrix *A*, degree matrix *D*, and combinatorial Laplacian *L* = *D* − *A*. For undirected graphs *L* is symmetric positive semidefinite with eigenvalues 0 = *µ*_1_ ≤ … ≤ *µ*_*n*_; the multiplicity of *µ*_1_ counts connected components, and *µ*_2_ (the algebraic connectivity) measures resistance to disconnection [109–111].

Linearising a network model gives a Jacobian whose *sparsity pattern* is dictated by topology: *J*_*ij*_ = 0 whenever *A*_*ij*_ = 0 for *i* ≠ *j*. Kinetics fill in admissible entries but cannot create new ones. This is the constraint exploited by the topological prior in (22).

For diffusively coupled systems with identical node dynamics,

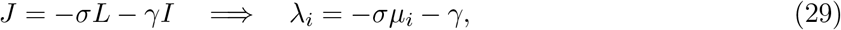

the cleanest instance of the correspondence.

### 9.2 Bounds Valid for Non-Symmetric Generators

#### Remark 9.1

(Weyl’s inequalities do not apply). It is sometimes asserted that for *J* = *J*_1_ + *J*_2_ with *J*_1_ topological and *J*_2_ kinetic, Weyl’s inequality bounds *λ*_*i*_(*J*) between *λ*_*i*_(*J*_1_)+*λ*_*n*_(*J*_2_) and *λ*_*i*_(*J*_1_)+*λ*_1_(*J*_2_). Weyl’s inequalities are a theorem about Hermitian matrices. Take 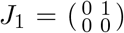 and 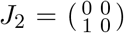: both are nilpotent with spectrum {0, 0}, so the putative bound collapses to [0, 0], yet *J*_1_ + *J*_2_ has spectrum {+1, −1}. For generically non-symmetric biological Jacobians with complex spectra the inequality is not merely inaccurate but ill-formed, since no canonical eigenvalue ordering exists. Three valid alternatives:

#### Gershgorin discs

Every eigenvalue of any *J* lies in

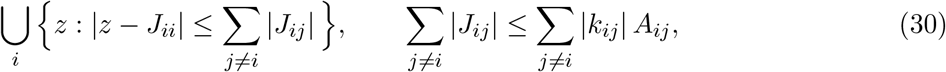

the right-hand bound depending only on topology and on bounds for kinetic constants.

#### Bendixson bounds

Decompose *J* = H + K with H = <u>^1^</u> (*J* + *J*^⊤^) and K = <u>^1^</u> (*J* − *J*^⊤^). Then

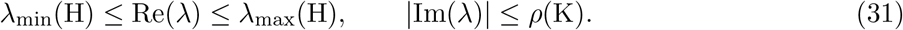

Weyl’s inequalities *do* apply to H, so the real-part bound inherits topological structure legitimately.

#### Bauer–Fike perturbation

Equation (11) bounds the departure from a nominal topological generator, at the cost of a factor *κ*(*V* ) quantifying non-normality.

### 9.3 Interlacing Under Structural Change

Since *L* is symmetric, Weyl’s inequalities are legitimate here.

#### Edge addition

Adding an edge adds a rank-one positive semidefinite term, so *µ*_*k*_(*G*) ≤ *µ*_*k*_(*G*+*e*) ≤ *µ*_*k*+1_(*G*).

#### Vertex deletion

The Laplacian of *G* − *v* is not a principal submatrix of *L*(*G*), since deleting a vertex also decrements neighbours’ degrees. Cauchy interlacing does not apply directly; the correct statement is Lotker’s weak interlacing theorem [112, 113]: with ascending ordering,

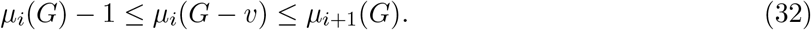

##### Remark 9.2.

The −1 cannot be dropped. For the path *P*_3_ with *µ* = (0, 1, 3), deleting the middle vertex leaves two isolated vertices with *µ* = (0, 0); the inequality *µ*_2_(*G*) ≤ *µ*_2_(*G* − *v*) would require 1 ≤ 0. The weakened form holds.

### 9.4 Controllability and Synchronizability

A network is defined as structurally controllable from a specific set of driver nodes if there exists at least one configuration of edge weights that renders the system controllable. Under this framework, the minimum number of driver nodes is determined by the maximum matching of the network [114], which constitutes a purely combinatorial property. For systems of identically coupled oscillators, the master stability function formalism [115] simplifies synchronization stability to a scalar condition evaluated at *σµ*_*k*_; consequently, overall synchronizability is dictated by the eigenratio *µ*_*n*_*/µ*_2_. Notably, small-world architectures are known to achieve minimized eigenratios, thereby enhancing synchronization stability [116].

### 9.5 Caveats

The aforementioned frameworks assume an underlying undirected, unsigned graph structure. However, biological processes such as gene regulation and intracellular signaling are inherently directional, necessitating the deployment of directed Laplacians characterized by complex spectra [117]. Furthermore, because activation and inhibition interactions carry opposing signs, a signed adjacency matrix fails to yield a positive semidefinite Laplacian [118], while system non-normality concurrently degrades eigenvalue conditioning (Section 3.6). While the analytical bounds detailed in Section 9.2 remain robust to all three structural complications, the expression in (29) does not; consequently, it should be restricted exclusively to systems where its underlying hypotheses have been explicitly verified.

### 9.6 When Topology Dominates

Under strong coupling, *σ* ≫ *γ* in (29) gives *λ*_*i*_ ≈ −*σµ*_*i*_: kinetics set only an overall scale. For pure consensus or diffusion the correspondence is exact. And when intra-module dynamics are fast relative to inter-module dynamics, nodes within a module equilibrate to a common value and the slow dynamics are governed by the Laplacian of the *quotient* graph. The precise conditions on the partition are established for consensus dynamics through external equitable partitions [119, 120], which is the natural formalisation of spectrum-preserving coarse-graining [121–123]; singular perturbation theory supplies the asymptotics [124]. At physiologically relevant observation timescales, the operator infers the quotient topology (the effective network as seen at that timescale) which is what makes the exported description compact.

## 10 Effective Topology Inference and the Cost of Assuming Isotropy

### 10.1 The Isotropic Result

For the diffusive case *J* = − *σL* − *γI* with symmetric *L and isotropic noise Q* = *qI*, the Lyapunov equation gives 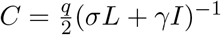 and hence

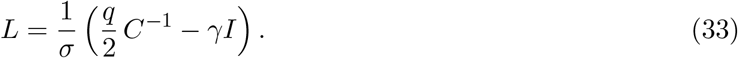

The precision matrix recovers the Laplacian up to scale and shift; its off-diagonal support coincides with the edge set. This correspondence is well established in graph signal processing [35–41] and in Gaussian graphical modelling [42], and is not claimed here.

### 10.2 What Happens When *Q* is Not Isotropic

Section 3 established that *Q* is generically anisotropic. Two questions must be separated, and conflating them is the common error: whether isotropy is *necessary* for (33) (it is not, and Proposition 10.2 gives the exact condition) and how badly the correspondence fails when that condition is violated, which is what Proposition 10.1 and Table 9 quantify.

**Table 9:** Recovery of the Laplacian from the precision matrix under three noise models. Under isotropic noise, recovery is exact and non-edges of *L* map to exact zeros of Θ. Under realistic anisotropy the fit collapses, and the largest phantom entry *exceeds* the average true edge, so a thresholding procedure returns predominantly false edges.

| Noise model | $R^2(\Theta_{\text{off}} \text{ vs } L_{\text{off}})$ | Spurious-edge leakage |
| --- | --- | --- |
| Isotropic, $Q = qI$ | 1.000 | 0.000 |
| Anisotropic diagonal, 5 decades of dynamic range | 0.257 | 2.247 |
| Low-rank extrinsic, $Q = gg^\top + 0.1I$ | 0.341 | 1.335 |

#### Proposition 10.1

(Covariance under general noise). *Let L* = *U* diag(*µ*_*i*_) *U*^⊤^ *and J* = −*σL* − *γI. Write* 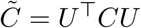 *and* 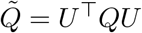. *Then*

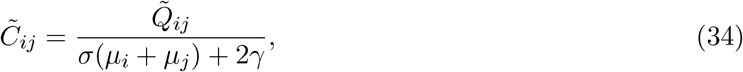

*i*.*e*. 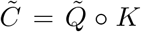 *with Cauchy kernel K*_*ij*_ = [*σ*(*µ*_*i*_ + *µ*_*j*_) + 2*γ*]^−1^ *and* ∘ *the Hadamard product. Consequently, if* 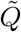 *has any nonzero off-diagonal entry then C*^−1^ *is not a function of L alone and the eigenvectors of L are not those of C*.

*Proof*. Substituting *J* = −*σL* − *γI* into (17) and conjugating by *U* gives 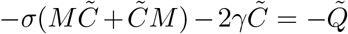 with *M* = diag(*µ*_*i*_); reading off entry (*i, j*) yields (34). If 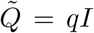 then 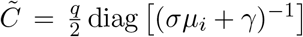 recovering (33); otherwise 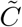 has off-diagonal entries in the Laplacian eigenbasis and its inverse is not a function of *L* alone. □

#### Proposition 10.2

(Exact condition for the affine correspondence). *Let J* = −*σL* − *γI with L symmetric, and let C solve* (17). *Then C*^−1^ = *aL* + *bI for scalars a, b with a* ≠ 0 *if and only if* [*Q, L*] = 0 *and the modal noise powers* 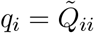 *satisfy*

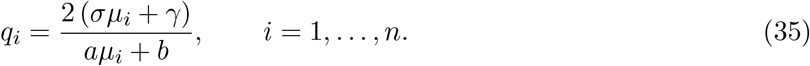

*Isotropy is the special case σ/a* = *γ/b. Condition* (35) *defines a genuinely anisotropic family: for any L with at least two distinct eigenvalues and any admissible* (*a, b*) *with σ/a* ≠ *γ/b, the q*_*i*_ *are non-constant*.

*Proof*. Commutation is necessary because *C*^−1^ = *aL* + *bI* shares its eigenbasis with *L*, and by (34) 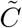 is diagonal only if 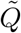 is. Given [*Q, L*] = 0, (34) gives 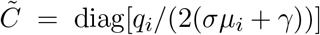, so *C*^−1^ = *U* diag[2(*σµ*_*i*_ + *γ*)*/q*_*i*_] *U*^⊤^, which equals *aL* + *bI* exactly when 2(*σµ*_*i*_ + *γ*)*/q*_*i*_ = *aµ*_*i*_ + *b* for every *i*. Solving for *q*_*i*_ gives (35); constancy of *q*_*i*_ in *µ*_*i*_ forces *σ/a* = *γ/b*. □

#### Remark 10.3

(The surviving family is not a measure-zero curiosity in *q*, but it is a strong condition on the pair (*Q, L*)). On the eight-node ring with two chords used below, with *σ* = 1, *γ* = 0.3: (*a, b*) = (1, 1) gives modal noise powers spanning a factor of 2.93 and (*a, b*) = (0.7, 2) a factor of 6.32, and in both cases ∥*C*^−1^ − (*aL* + *bI*)∥_max_ is 1.3 × 10^−15^ relative to ∥*aL* + *bI*∥_max_, about 6 *ε*_mach_ (*ε*_mach_ ≈ 2.2 × 10^−16^), i.e. at the level of double-precision rounding error. So a two- to sixfold spread in modal noise power is entirely compatible with exact Laplacian recovery, and reporting anisotropy alone is not grounds for rejecting a precision-matrix estimate.

What the condition demands is not near-isotropy but *alignment* : *Q* must commute with *L*, and its modal powers must track (35) along the Laplacian spectrum. Neither is implied by the stoichiometric form (4), and there is no mechanism that would enforce them, which is why the generic case below is the relevant one for practice. The correct operational statement is therefore conditional rather than prohibitive: a precision-matrix estimate requires an explicit and justified claim about the relation between the driving noise and the network, and an appeal to sparsity is not such a claim.

Even commutation alone does not rescue the weaker practice of reading module structure off the covariance. Under [*Q, L*] = 0 the covariance eigenvalues are the modal variances 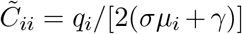, so ranking modes by variance reproduces the ordering of the Laplacian eigenvalues exactly under isotropy (Spearman *ρ* = −1 whenever the spectrum has no ties). Under commuting anisotropy that ordering degrades monotonically in the spread of *q*. We drew 25 connected Erdős–Rényi graphs on eight nodes with edge probability 0.3, keeping only graphs whose Laplacian eigenvalues are separated by more than 10^−6^, and set *σ* = 1, *γ* = 0.3. For each graph we drew 30 modal noise vectors with log_10_ *q*_*i*_ independent and uniform on [0, *k*], so that *k* is the spread in decades, and computed *ρ* between modal variance and *µ*_*i*_ over all eight modes. Across the resulting 750 graph–draw pairs (mean ± standard deviation), *ρ* = −0.92 ± 0.06 at half a decade, −0.77 ± 0.14 at one decade, −0.54 ± 0.26 at two and −0.38 ± 0.32 at three. Transcriptional noise power spans four to five decades [56, 57].

We now quantify the generic case numerically. On the same eight-node ring with two chords, with *σ* = 1, *γ* = 0.3, we compute *C* under three noise models, invert it, and regress the off-diagonal entries of Θ = *C*^−1^ on those of *L*. Table 9 reports the coefficient of determination and the *spurious-edge leakage*: the largest magnitude among entries of Θ at non-edges, divided by the mean magnitude at true edges.

#### Interpretation.

Network inference by thresholding a precision matrix requires condition (35), of which isotropy is one case. Nothing in the stoichiometric structure of biochemical noise enforces either half of it, and where it fails the failure does not merely add noise to the estimate. It creates strong apparent edges where none exist, in the tabulated cases outnumbering the true ones. The remedy is not a better threshold. It is to estimate *Q* (Section 5.5) and then solve (34) for *L*, which is a well-posed linear problem once 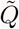 is known, and, failing that, to report (35) as an assumption rather than to leave it implicit.

### 10.3 Effective versus Structural Topology

Two notions of network must be distinguished, and neuroscience has maintained the distinction for three decades under the terms we adopt [33, 34]. *Structural topology* is the network of possible interactions: binding partners, regulatory relationships, spatial adjacency. *Effective topology* is the network of dynamical influence as inferred from fluctuations, i.e. not which interactions exist but which are active under current conditions.

The operator recovers effective topology. Discrepancies with structural topology are not necessarily errors: a structural edge may be kinetically silent, an indirect path may dominate a direct one, and feedback may generate apparent coupling between structurally distant nodes.

### 10.4 Diagnostics

| Observed pattern | Interpretation |
| --- | --- |
| $\hat{Q}$ from (20) not positive semidefinite | Coloured noise, hidden variables, non-stationarity, or failed linearisation (Protocol 5.7) |
| $\hat{R}(\tau)$ smooth at $\tau = 0$ | Driving noise is coloured; augment per (7) (Protocol 5.8) |
| Slow modes with eigenvectors delocalised over unrelated species | Extrinsic modulator mistaken for network mode (Remark 3.4); regress out covariates |
| $\hat{C}_{cs} \gg \hat{C}_{long}$ | Population heterogeneity; regime H of Table 6 |
| $EB(T)$ not decaying with $T$ | Ergodicity breaking; regime N |
| Inferred mixing faster than topology predicts | Structural network incomplete |
| Inferred mixing slower than predicted | Structural edges kinetically inactive |
| Effective rank exceeds structural rank | Hidden variables coupling to observed nodes |
| Large $\text{dep}_F(\hat{J})$ or $\kappa(\hat{Y})$ with unstable eigenvalue estimates | Strong non-normality; eigenvalue-based specification unreliable |

### 10.5 Information Fusion

Topological priors and dynamical evidence should be combined rather than compared. Place

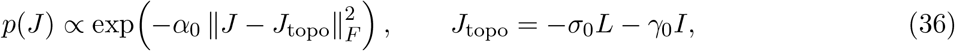

and combine with the likelihood implied by (22); equivalently, the penalty 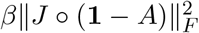 enforces the structural support directly. This follows [29], who demonstrated for metabolic networks both the topological constraint on *J* and the derivation of a non-diagonal noise structure from stoichiometry. Their finding that correct noise structure improves precision at the cost of recall is directly relevant: *Q* is not a nuisance parameter but a determinant of what the inference returns.

## 11 Canonical Circuits

### 11.1 Secrete-and-Grow: Compensation and Spectral Collapse

The endocrine secrete-and-grow circuit [98, 105] is

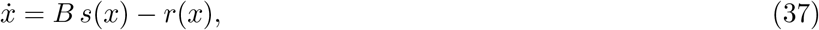

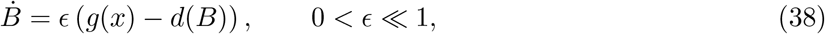

with *x* a hormone level and *B* a gland mass. The Jacobian carries a fast eigenvalue of order unity and a slow one of order *ϵ*.

Note that *Q* here is emphatically not isotropic: hormone turnover *s, r* operates at order unity while gland turnover *g, d* operates at order *ϵ*, so by (4) the modal noise powers differ by the same factor. By (10) the two effects (slower damping and weaker forcing) partially cancel, and which mode dominates the covariance depends on the ratio, not on the timescale separation alone. This is the simplest concrete illustration of Section 3.5.

The circuit exhibits dynamical compensation: multiple parameter configurations (*s, g, d*) collapse onto the same observable mode, and the operator correctly returns a one-dimensional description with high uncertainty orthogonal to it.

### 11.2 Bistable Fibrosis: Spectral Bifurcation

Autocrine–paracrine circuits coupling macrophages and myofibroblasts form a bistable system with healthy and fibrotic attractors separated by a saddle [125]. Approaching the saddle, one eigenvalue tends to zero from below, which is critical slowing down [126–128]. The eigenvalue trajectory defines a therapeutic window. Section 12.3 treats the reliability of that inference.

Here the ergodicity question is acute. Cross-sectional tissue data mix cells from both basins, so the population covariance is a mixture and Theorem 4.1 applies with a bimodal *π*. The inferred slow eigenvalue is then biased toward zero for two independent reasons (heterogeneity inflation and genuine proximity to bifurcation) which cannot be separated without either stratification by basin or longitudinal data.

### 11.3 Two-Gland Oscillator

Endocrine axes such as the hypothalamic–pituitary–adrenal system behave as coupled-gland oscillators [129–131]. A linearised representation is

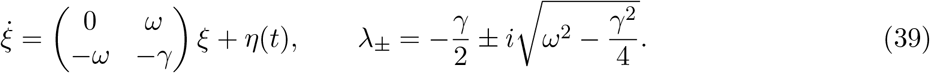

With isotropic noise *Q* = *qI* the Lyapunov equation gives

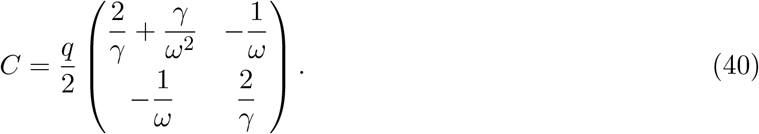

The sign of the off-diagonal entry is negative and is not conventional: it encodes the phase relationship, and reversing it reverses the inferred lead–lag structure. For *ω* = 2, *γ* = 0.5, *q* = 1 this gives *C*_11_ = 2.0625, *C*_12_ = −0.25, *C*_22_ = 2.

By Corollary 5.3 this circuit is exactly where equal-time data are insufficient: Table 7 was constructed from (39) and shows that *ω* is entirely undetermined by *C* alone. Oscillatory endocrine circuits are therefore the strongest argument for the lag-based or velocity-based definition of Section 6.

## 12 Non-Stationarity and Time-Varying Inference

Stationarity represents perhaps the most consistently violated assumption across biological systems. For instance, endogenous circadian rhythms continuously modulate physiological parameters throughout a 24-hour cycle, while developmental processes fundamentally transform system dynamics over multi-week intervals. Furthermore, disease progression drives systematic shifts in operational regimes, and clinical or experimental interventions are deliberately introduced to perturb the system’s baseline operating point.

### 12.1 Quasi-Stationary Windows

If parameters vary slowly relative to relaxation, the system tracks an evolving equilibrium and satisfies an instantaneous Lyapunov equation. The operator may then be applied within sliding windows of width *W* satisfying

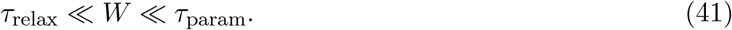

Combined with (28) this is a demanding double inequality: the window must contain enough effectively independent samples to estimate an *n*×*n* covariance while remaining short compared with drift. When the two conflict, no window width is valid and dimension must be reduced before windowing.

Windowing strategies (rectangular, exponentially weighted, Gaussian-kernel) trade time resolution against estimation variance in the standard way. Sliding covariance tracking is the established approach in dynamic functional connectivity [132, 133], where its artefacts are well documented.

### 12.2 Regime Detection

Where change is discrete rather than continuous, the record must be segmented before stationary analysis; mixing regimes produces a covariance describing no actual operating condition. Change-point detection may use CUSUM, Bayesian online detection [134], or direct monitoring of spectral change.

#### Attribution.

Frameworks incorporating a latent discrete regime to index a collection of linear Gaussian dynamical systems (enabling the joint inference of time-series segmentation and regime-specific dynamics) are formally classified as switching linear dynamical systems (SLDS), a paradigm supported by a mature inferential literature [135–139]. Recurrent extensions of these models condition regime transitions directly upon the continuous state variable, which is particularly appropriate when such transitions are driven by underlying physiological mechanisms. Consequently, future implementations should leverage these established methodologies rather than attempting de novo derivations.

### 12.3 Critical Slowing Down and Early Warning

As a dynamical system approaches a saddle-node bifurcation, it undergoes critical slowing down: its dominant eigenvalue converges toward zero, the relaxation time diverges, and both variance and auto-correlation exhibit significant growth. This phenomenon is well established across multiple disciplines. The utilization of the slowest decay rate as a precursor originated with the concept of degenerate fingerprinting [140], while a substantial body of ecological literature subsequently formalized these statistical indicators [126, 127, 141, 142]. In biomedical contexts, this principle has been instantiated as the dynamical network biomarker framework [143, 144]. Furthermore, multivariate indicators utilizing covariance eigendecomposition have been thoroughly benchmarked [145, 146], and empirical Jacobian reconstruction has recently been integrated directly with early warning systems [147].

#### The correct extrapolation law

Near a saddle-node with control parameter 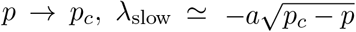, so under linear parameter drift the eigenvalue is not linear in time.

##### Proposition 12.1

(Linear extrapolation overestimates by a factor of two). *With λ*(*t*) = −*a*^*′*^(*t*_*c*_ −*t*)^1*/*2^, *linear extrapolation of λ to zero predicts a remaining time of* 2(*t*_*c*_ − *t*): *exactly twice the true value*.

*Proof*. 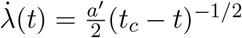, so 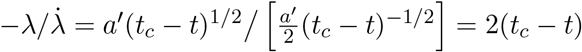. □

The error reports twice the available intervention window. The correct procedure fits 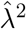, which is linear in *t*.

#### Reliability caveats

Statistical power remains limited and false negatives are frequently encountered at realistic noise levels [148]; consequently, the absence of an early warning signal cannot be construed as evidence of system stability. More specifically, a systematic evaluation of covariance-based network early-warning indicators demonstrated that off-diagonal covariance entries failed to improve detection quality relative to the variances of well-chosen sentinel nodes [149]. While we consider this an open empirical question, we note that it does not preclude the utility of the operator for reduced-order control design, where mode shapes are required a priori. Nonetheless, we recommend benchmarking new metrics against node-variance baselines rather than assuming intrinsic methodological superiority. Finally, Corollary 4.2 introduces a distinct analytical hazard: cross-sectional data independently bias *λ*_slow_ toward zero, meaning that population heterogeneity can erroneously masquerade as a proximal critical transition.

### 12.4 Transient Response Analysis

Where the system is perturbed, transients yield spectral information without stationarity. Following an impulse, 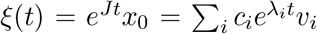 with 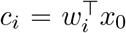, so fitting a sum of exponentials by Prony’s method, matrix pencil, or ESPRIT [150, 151] recovers eigenvalues directly. The coefficients depend on the *left* eigenvectors, which is the formal reason actuator placement is governed by *W*_*r*_ while readout interpretation is governed by *V*_*r*_ — a statement that presupposes the eigen-refinement of Remark 6.3 and should be qualified where it does not apply. Choosing *x*_0_ = arg max_∥*x*∥=1_ min_*i*_ |*c*_*i*_(*x*)| maximises the weakest mode excitation and is the natural objective for experimental design [152].

### 12.5 Periodic Modulation

For circadian systems the generator is periodic and stability is governed by Floquet exponents; Floquet analysis of circadian oscillators is standard [153]. Phase-resolved analysis pools data by circadian phase, yielding a spectral map around the cycle and connecting to chronotherapy [154, 155].

#### Remark 12.2.

The relation 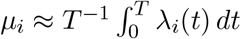 between Floquet exponents and time-averaged instantaneous eigenvalues holds only adiabatically. There exist periodic systems in which *J*(*t*) is Hurwitz at every instant yet the Floquet system is unstable. Since circadian modulation is not slow relative to many signalling timescales, the adiabatic condition must be checked.

### 12.6 Algorithm

#### Algorithm 1

UNS with noise recovery, ergodicity screening and regime detection

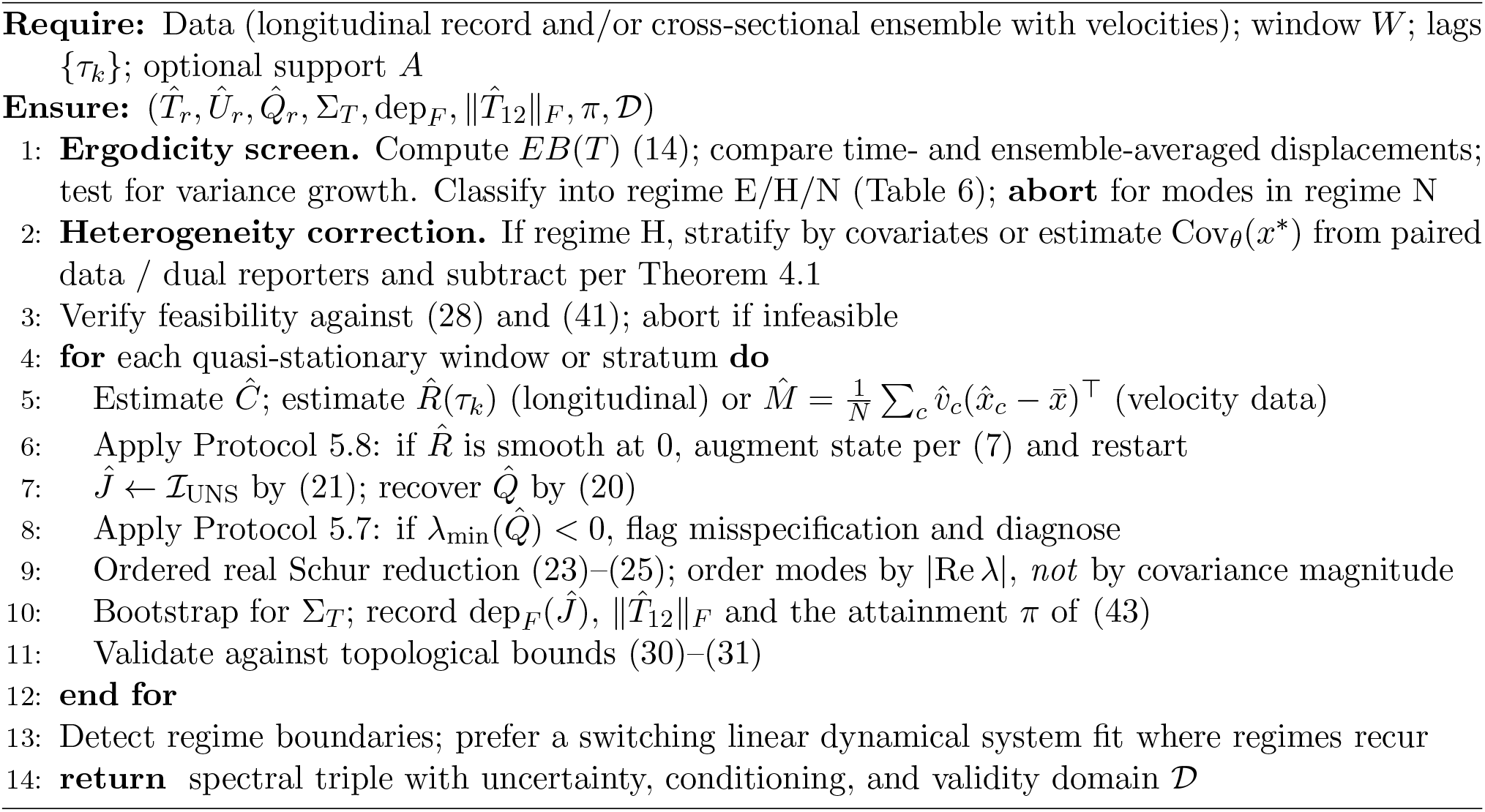

## 13 Nonlinear Extensions

### Failure modes

*Rank collapse* from dynamical compensation, diagnosed by rapid singular-value decay in *Ĉ* or near-singularity of the Fisher information. *Critical slowing*, where the quasi-Gaussian assumption degrades and autocorrelation decays non-exponentially. *Regime switching*, diagnosed by significant change-point statistics.

### Piecewise-linear representation

For systems with a small number of physiologically meaningful operating modes, partition the state space and fit a generator per region as in Section 12.2, retaining the identifiability analysis and uncertainty machinery within each.

### Koopman lifting

For globally nonlinear behaviour, lift to a space of observables where the dynamics are approximately linear [156–160]. A productive combination uses the local reduction to identify modal coordinates near the primary equilibrium and builds the lift around them.

### Higher-order statistics

Third and fourth cumulants carry information about nonlinear interaction that covariance misses, and tensor decompositions [161] can extract modal structure from them. The sample requirements are severe (already binding at second order by (28)) so we record this as a direction rather than a capability.

## 14 The Spectral Triple as an Interface Contract

### 14.1 What Crosses a Scale Boundary

In a hierarchical modelling architecture, a component at one scale must export a compressed description upward. It should not export raw states (too numerous, violating R2) or mechanistic parameters (unidentifiable, violating R1). We propose that it export

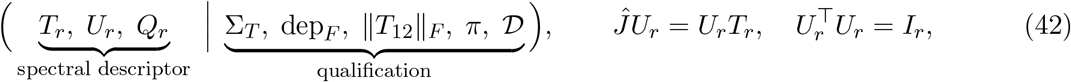

where the descriptor is the description proper and the second group qualifies it. *T*_*r*_ specifies timescales, frequencies and stability margins through spec(*T*_*r*_); *U*_*r*_ specifies which combinations of the underlying variables the description is about; *Q*_*r*_ specifies modal fluctuation amplitudes, *recovered rather than assumed*. Where the eigen-refinement of Remark 6.3 is available it is exported alongside, since *V*_*r*_ carries the observable mode shapes and *W*_*r*_ the directions along which the modes are excited (which is why actuator placement is governed by *W*_*r*_ and readout interpretation by *V*_*r*_) together with *κ*(*Y* ); where it is not available, that fact is itself part of *D*.

Of the qualifiers, Σ_*T*_ is the uncertainty on the reduced generator (reported on spec(*T*_*r*_) and on col(*U*_*r*_) as principal angles, both being invariant under the basis freedom of Remark 6.5); dep_*F*_ and ∥*T*_12_∥_*F*_ quantify departure from normality and coupling from the discarded block (Remarks 6.6 and 6.4), and hence how far the spectral description can be trusted and whether it may be read as a subsystem model at all; *π* is the attainment of Remark 14.1; and *D* declares the validity domain: operating regime, stationarity window, ergodic regime per Table 6, the conditioning of *Ĉ* per Section 7.3, and the explicit list of directions found unidentifiable.

The qualification group is not bookkeeping. A contract exporting a point estimate without its uncertainty and without its domain of validity is not a contract but an assertion. In particular, D must record whether the estimate came from ergodic longitudinal data, from a stratified cross-sectional ensemble, or from velocity statistics, since the failure modes differ and downstream consumers must know which apply.

#### Remark 14.1

(Attainment: why uncertainty is not sufficient qualification). A descriptor can satisfy all of the aforementioned criteria and yet remain fundamentally misleading, because Σ_*T*_ quantifies estimation precision rather than the proportion of the underlying system dynamics recovered by the estimator. Crucially, these two metrics diverge in a problematic direction: a highly precise descriptor that captures negligible information regarding the true system will be reported as a high-confidence result, with neither its uncertainty bounds nor its validity domain flagging the structural deficiency.

The missing quantity is two-sided. Let *e*_obs_ be the agreement between the exported description and a held-out observable, on any fixed scale. Let *e*_∅_ be the same quantity for a null that preserves everything about the design except the structure being claimed (for a network-supported generator, a degree-preserving rewiring; for a modal claim, a phase randomisation) and let *e*_⋆_ be a *design-matched ceiling* : the same quantity when a generator of the assumed class is planted in the true system and passed through the identical estimation pipeline at the identical sample size. Then

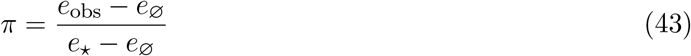

places the result on the interval the design actually admits. *π* ≈ 1 means the description recovered what was recoverable; *π* ≈ 0 means it did not, whatever Σ_*T*_ says. Reporting *e*_obs_ against *e*_∅_ alone is the common error and is not sufficient: a statistically significant improvement over a null can still be a small fraction of the attainable range, and without *e*_⋆_ there is no way to tell whether the shortfall lies in the data, the estimator, or the model class.

Two cautions follow from the construction. A narrow interval *e*_⋆_ − *e*_∅_ inflates *π* at small absolute lift, so the absolute lift must be reported alongside the ratio. And a *π* without an explicit *e*_∅_ test is not a result: a structure-preserving null that the estimate fails to beat means the apparent attainment is an artefact of whatever the null preserves.

### 14.2 Bidirectional Consistency

The structural contract imposes consistency constraints in both directions. *Upward* : any coarser, macro-scale model must remain strictly consistent with the exported descriptors. For instance, if the macro-model predicts an oscillation with period *T*, then spec(*T*_*r*_) must contain eigenvalues satisfying Im(*λ*) = 2*π/T* within the confidence bounds defined by Σ_*T*_ . A coarse model that fails this reconciliation is rejected rather than tuned. *Downward* : any micro-scale refinement must faithfully reproduce these properties. Specifically, a more detailed model evaluated under the identical observation protocol must return the same operational triple within the established uncertainty. This bidirectional consistency imposes a powerful constraint on mechanistic elaboration, effectively mitigating the parameter proliferation that traditionally compromises forward multiscale modeling.

Bidirectionality distinguishes this from ordinary model reduction: a reduction produces a smaller model, whereas a contract produces an obligation.

### 14.3 Relation to Contract Theory

The construction has a precise formal analogue. In cyber-physical systems design, an assume– guarantee contract specifies what a component requires of its environment and guarantees in return, with composition and refinement operators determining when contracts may be combined [162–164]. In control, the theory of hierarchically consistent abstraction determines when a controller synthesised on an abstract model refines to the concrete system [165–168].

Equation (42) is an assume–guarantee contract whose guarantee is spectral. The feature distinguishing the biological setting, and the reason the existing theory cannot be applied unmodified, is that the contract terms must be *inferred from data* rather than specified by a designer. Everything in Sections 3–7 exists to make that inference well posed, and identifiability is treated so carefully there because an unidentifiable contract term is a contract that cannot be checked.

### 14.4 Controllability Inheritance

#### Definition 14.2

(Controllability inheritance). Let 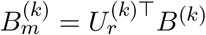 be the reduced input matrix at level *k* (or 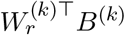 where the eigen-refinement of Remark 6.3 is in use; the rank condition below is the same in either basis). An abstraction from level *k* to *k* + 1 satisfies controllability inheritance if

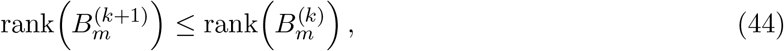

with equality on all modes designated as control targets.

If a coarse-graining loses controllability of modes controllable below, the abstraction is too aggressive: it has discarded degrees of freedom that intervention requires. Condition (44) therefore *falsifies* a proposed abstraction rather than merely scoring it, and it is checkable from quantities the contract already exports. The theorem underlying the corresponding statement for control systems is due to Pappas, Lafferriere and Sastry [165]; posing it as a design constraint on biological coarse-graining is, so far as we are aware, new.

A companion condition constrains spectral ordering: an abstraction should map eigenvalues monotonically, since an inversion of the ordering of timescales indicates that distinct modes have been improperly mixed. The formal machinery is that of equitable partitions and spectral coarse-graining [119–122].

### 14.5 Downstream Use

The exported triple is exactly what control synthesis consumes. The reduced state 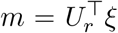 is the natural object for estimation and feedback; spec(*T*_*r*_) fixes sampling rate and stability requirements; *Q*_*r*_ tunes the Kalman filter; and where input data are available the modal input matrix 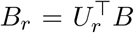 completes a state-space model on which linear-quadratic regulation, model predictive control, H_∞_ synthesis and Kalman filtering all operate directly [169, 170]. The uncertainty Σ_*T*_, the conditioning measures dep_*F*_ and ∥*T*_12_∥_*F*_, and the attainment *π* pass forward into robust synthesis, so a poorly identified (or a well-identified but uninformative) mode produces a conservative controller rather than a confident wrong one.

When the reduced-order description is employed to constrain a learned latent representation, the underlying mechanism takes the form of spectral regularization of the induced latent dynamics. While this constitutes a well-established computational technique [171–175], the key distinction in the present framework is that the regularization target is directly derived from empirical data rather than selected heuristically or manually predefined.

## 15 Limitations

Stating limitations precisely is part of the method, since (42) is only as good its validity domain.

### L1. Equal-time data are insufficient

By Theorem 5.2 the spectrum is not identifiable from (*C, Q*) alone. Implementations restricted to equal-time covariance (which includes much published metabolomics work, where sampling is destructive) recover only gauge-invariant quantities. This is a hard limit, not a numerical difficulty, and it applies to Definition 6.1 itself: at *α*_1_ = *α*_2_ = 0 the operator returns the gauge family of Theorem 5.2, so a study running that corner is not running a weakened version of UNS but a differently-posed problem, and should report the attainment *π* of Remark 14.1 rather than a point estimate of the spectrum.

### L2. Cross-sectional data are bias-limited

By Theorem 4.1 and Corollary 4.2, population heterogeneity biases relaxation rates downward, and the bias does not shrink with sample size. Stratification, a paired longitudinal panel, or dual-reporter designs are required.

### L3. Non-ergodic modes are excluded, not estimated

Slow epigenetic and glassy processes have no stationary distribution over feasible observation windows. The operator must declare them outside its domain rather than return wide error bars around a quantity that does not exist.

### L4. Isotropy is a hypothesis, not a default

By Proposition 3.1 and Remark 3.2 the diffusion matrix is generically anisotropic, though exact isotropy is attainable; by Section 3.5 anisotropy can invert mode ranking; and by Proposition 10.2 the precision-matrix correspondence survives under a condition that is weaker than isotropy but is still a substantive claim about the relation between *Q* and *L*, failing which Table 9 applies. Methods inherited from graph signal processing must not be transplanted without stating which condition is being assumed and why.

### L5. Non-normality limits what the spectrum specifies

Where dep_*F*_ (*Ĵ*) or *κ*(*Y* ) is large, eigenvalues are ill-conditioned by (11) and do not bound transient growth; and where ∥*T*_12_∥_*F*_ is comparable to ∥*T*_*r*_∥_*F*_ the descriptor is a restriction rather than a subsystem model (Remark 6.4). Pseudospectral analysis is the appropriate refinement [64, 70].

### L6. Colored extrinsic noise requires augmentation

The white-noise Lyapunov equation is invalid when modulators fluctuate on comparable timescales. Protocol 5.8 detects this; (7) corrects it, at the cost of introducing unobserved states.

### L7. Sample complexity binds

By (28), longitudinal resolution of an *n*-dimensional covariance requires observation over many multiples of the slowest timescale. This should gate study design, not be discovered afterwards.

### L8. Early-warning claims require benchmarking

Detection power is limited [148], and covariance-based network indicators have been found not to outperform sentinel-node variances [149].

### L9. Topological bounds require symmetric structure

The diffusive correspondence assumes undirected, unsigned coupling; for directed and signed networks only the weaker bounds of Section 9.2 apply [117, 118].

### L10. The covariance estimate is usually singular

Section 7.3: rank *Ĉ* ≤ min(*N* −1, *n*), so the standing hypothesis *Ĉ* ≻ 0 fails for the assay shapes the operator is aimed at. Definition 6.1 survives this because it never inverts *Ĉ*; the closed-form routes do not, and the naive repairs fail in opposite directions depending on whether the deficiency is structural or from sampling. Where *N* ≲ 2*n* the identified subspace is at most (*N* − 1)-dimensional whatever dynamical information is supplied, and *r* must be chosen well inside it.

### L11. The exported descriptor is a restriction, not a subsystem model

By Remark 6.4 the retained coordinates are driven by the discarded ones through *T*_12_, and closure requires a realisation of the discarded block rather than a summary of it. Downstream models that treat the descriptor as an autonomous subsystem are making an assumption that ∥*T*_12_∥_*F*_ is reported in order to expose.

## 16 Conclusion

Rational intervention requires knowing how a system responds to perturbation, and that knowledge resides in the local generator of the dynamics. Forward mechanistic models contain the generator but cannot be calibrated to recover it. Learned representations can be fitted but do not contain it. Unified Network Spectroscopy supplies the missing inverse step by exploiting the relationship between fluctuation and dissipation: spontaneous fluctuations about a physiological setpoint constitute a passive interrogation of the dynamics, weighted by (9) toward the modes that intervention must address.

Three qualifications, developed at length above, determine whether that interrogation succeeds. The driving noise is generically anisotropic. Its structure is fixed by stoichiometry as *Q* = Ω^−1^*S* diag(*a*)*S*^⊤^, generically non-diagonal with entries of both signs, spanning the dynamic range of the propensities and rank-limited by the reaction count; extrinsic modulation adds a low-rank, coloured component whose augmented generator is non-normal by construction and whose relaxation rates contaminate the inferred spectrum. Exact isotropy is attainable but sits on a null set of admissible flux vectors, so assuming it is not conservative: it can invert the ranking of modes, and outside the exact condition of Proposition 10.2 it destroys precision-matrix topology recovery so thoroughly that phantom edges exceed true ones. That condition is the useful form of the result, because it is checkable: it tells a practitioner what would have to hold, rather than only that isotropy will not.

The equal-time covariance does not identify the generator, nor even its spectrum. The Lyapunov inverse admits an *n*(*n* − 1)*/*2-dimensional gauge freedom whose parameter is the probability current, and living systems (sustained by dissipation in non-equilibrium steady states) sit generically off the identifiable submanifold. The obstruction has a compact reading: by (1) the equal-time covariance is the frequency-integrated power spectrum, so inverting it alone amounts to inferring a source’s line structure from its total brightness, i.e. photometry rather than spectroscopy. Lagged covariance restores the frequency resolution, and so does a per-cell velocity estimate, with the useful consequence that the diffusion matrix becomes a recovered and falsifiable quantity rather than an assumption.

Ensemble statistics substitute for temporal statistics only under an ergodic hypothesis with three separate conditions, of which population homogeneity is the one biology violates. The resulting bias is systematic and, unlike a variance, does not diminish with sample size; it makes systems appear closer to bifurcation than they are. Slow epigenetic modes fail ergodicity outright and must be excluded from the description rather than estimated poorly.

None of the underlying machinery is new. The Lyapunov inversion has been developed independently in metabolomics, in computational neuroscience and in graph signal processing, and Section 2 sets out that inheritance. What is new is the quantification of how the standard isotropy and ergodicity assumptions fail for biological data, the constructive remedies for each, and the use of the resulting spectral triple as a typed interface contract — the commitment that this and nothing else crosses a scale boundary, carrying its uncertainty, its conditioning and its validity domain, with obligations enforced in both directions and with controllability inheritance falsifying any abstraction that has compressed away the capacity to intervene.

That is a modest claim by the standards of methodological novelty and a substantial one by the standards of practice. The components have existed for some time. What has been missing is a discipline for composing them such that each scale exports only what the data support, each interface can be checked, and each abstraction can be falsified.

